# Targeting CPSF73, the mRNA 3’ End Processing Endonuclease, Moves Cancer Cells Away from the Mesenchymal State

**DOI:** 10.64898/2026.03.01.708894

**Authors:** Marzieh Naseri, Huiyun Liu, Luyang Wang, Salwa Mohd Mostafa, Ehsan Ranaei Pirmardan, Bin Tian, Claire L. Moore

**Affiliations:** Department of Developmental, Molecular, and Chemical Biology, Tufts University School of Medicine, Boston, MA 02111, USA; The Wistar Institute, Philadelphia, PA, 19104, USA; Graduate School of Biomedical Sciences, Tufts University School of Medicine, Boston, MA 02111, USA

**Author notes:** **Corresponding author:** Claire Moore, Ph.D., Department of Developmental, Molecular, and Chemical Biology, Tufts University School of Medicine, Jaharis Bldg - 150 Harrison Ave, Boston, MA 02111, USA.

**Keywords:** CPSF73, Alternative polyadenylation (APA), Epithelial-mesenchymal transition (EMT), JTE-607, 3′ UTR lengthening

## Abstract

**Background:** Metastasis significantly contributes to cancer-related mortality and therapeutic failure. Cancer cells acquire metastatic potential by losing epithelial characteristics and gaining mesenchymal properties through the epithelial-mesenchymal transition (EMT). Differential poly(A) site (PAS) usage, known as alternative polyadenylation (APA), generates mRNA isoforms differing in coding sequence, subcellular localization, stability, or translation efficiency. In cancer, 3′UTR shortening increases expression of proto-oncogenes by escaping miRNA-mediated repression. High expression of CPSF73, which cleaves mRNA precursors at PASs, is associated with unfavorable prognoses in cancer patients. However, the role of APA in regulating EMT remains poorly understood.

**Methods:** In this study, to investigate the role of APA in EMT, we employed JTE-607, a small-molecule inhibitor of CPSF73 activity, to examine the impact of catalytic inhibition of CPSF73 on proliferation and EMT in MDA-MB-231, MCF7, A549, and HepG2 cancer cells. To identify differential usage of PASs, global profiling of APA changes, and differential gene expression analysis were performed in MDA-MB-231 cells. Additionally, antisense oligonucleotides were used to block the use of a specific PAS whose APA change may be a driver of EMT reversal.

**Results:** Our findings showed that catalytic inhibition of CPSF73 not only attenuates cancer cell proliferation but also moves the cells away from the mesenchymal state across all four cell lines tested. Global profiling of APA changes following CPSF73 inhibition revealed widespread 3′UTR lengthening and suppression of intronic PASs in MDA-MB-231 cells. APA shifts were observed in key EMT-related genes, accompanied by decreased expression of corresponding proteins across all four cell lines. We used antisense morpholino oligonucleotides to block the proximal PAS of AKT2, shifting the balance of AKT2 mRNA isoforms toward the long isoform. This shift caused EMT reversal, marked by reduced AKT2 protein expression, changes in EMT-related markers, and impaired invasion by MDA-MB-231 cells.

**Conclusion:** Together, these findings identify APA-mediated 3’UTR lengthening, with functional consequences in EMT-related genes, as a coordinated mechanism leading to an attenuated EMT phenotype, highlighting a significant connection between APA and the EMT process. Interfering with these APA changes may offer a promising therapeutic strategy to suppress metastasis, with potential efficacy across multiple pathways.

**Statement of Significance:** Our findings highlight APA-mediated 3’ UTR lengthening as a coordinated mechanism that promotes EMT reversal and support CPSF73 inhibition or APA-targeting strategies as potential therapeutic approaches to suppress metastasis across multiple pathways.

## Introduction

Metastasis is a major contributor to cancer-related deaths and often leads to the failure of conventional cancer therapeutic strategies (1–4). Cancer cells acquire aggressive characteristics through several mechanisms, including the epithelial-mesenchymal transition (EMT). EMT is a process by which epithelial cells lose their defining epithelial characteristics, such as cell-cell adhesion and polarity, and acquire properties of mesenchymal cells that enhance cell motility (5, 6). This transition to a more mesenchymal state is associated with alterations of transcriptional programs that modify the expression of adhesion molecules, resulting in increased metastasis capacity and therapeutic resistance of the tumor cells (7, 8). Reprogramming of gene expression by EMT transcription factors (EMT-TFs) such as Twist1, Zeb1, Zeb2, Slug, and Snail represses E-cadherin, an invasion suppressor gene, and activates dynamic changes in cellular organization. These changes, in turn, confer motile and invasive behaviors to cancer cells marked by increased expression of N-cadherin (9, 10). However, despite existing knowledge about the EMT process, significant challenges to developing new therapeutics remain due to the incomplete understanding of the mechanisms governing EMT and metastasis. Hence, identifying the molecular mechanisms that regulate the EMT-driven pathways and finding strategies to inhibit them is crucial to deciphering the drivers of cancer invasion and metastasis, ultimately leading to improved treatment outcomes.

One mechanism of regulating gene expression involves changes in cleavage and polyadenylation (C/P) of nascent RNA (pre-RNAs). The cleavage of the mRNA precursor at the 3’ end creates a site known as the polyadenylation site (PAS) for adding a chain of adenosines, called the poly(A) tail. This process also triggers transcription termination, thus preventing interference with downstream gene expression. The poly(A) tail protects mRNA from degradation, aids in its nuclear export to the cytoplasm, and facilitates its translation (11, 12).

C/P also provides a pervasive mechanism of post-transcriptional regulation in both physiological and pathological conditions (13–15). Altering the position of the PAS, resulting in alternative polyadenylation (APA), produces mRNA isoforms with different lengths of coding sequence or 3′ untranslated region (UTR). Shortening the 3′ UTR removes binding sites for regulatory elements such as microRNAs and non-coding RNAs, thereby influencing RNA stability, translation, and localization, whereas coding region shortening can dramatically alter protein function (16). Since nearly 70% of human genes have multiple PASs, APA significantly contributes to transcriptome diversity (17, 18).

The profile of APA site usage undergoes substantial changes during processes such as cell differentiation, development, and cancer (19–21). Dysregulation of C/P and dynamic APA events during tumorigenesis can promote cancer initiation and drive tumor metastasis (15, 22, 23). Widespread 3′ UTR shortening has been observed across multiple cancer types, and often causes increased mRNA expression that leads to the activation of transcription factors and proto-oncogenes by escaping miRNA- or RBP-mediated repression (21, 24, 25). 3’ UTR shortening has been reported to regulate the expression of some key driver genes in cancer progression and EMT, such as ZEB1, the multidrug resistance protein MRP-5/ABCC1, TWIST, and Aurora Kinase A (26–29). Additionally, Transforming Growth Factor (TGF)-β-mediated induction of EMT in human mammary epithelial cells caused APA changes of several EMT-relevant mRNAs regulated by the RNA-binding protein Quaking (QKI) (30). While these findings support an important regulatory role for APA in the expression of EMT-related genes, the possibility of manipulating APA changes as a novel strategy for therapeutic intervention in cancers to reverse EMT has not been rigorously explored.

CPSF73 (also called CPSF3), the nuclease that cleaves RNA at the PAS, has received considerable attention as a target for cancer therapy because high levels correlate with poor patient survival (31, 32), and as an enzyme, it is more readily druggable than other subunits of the C/P complex that serve RNA-binding or scaffolding functions (33). Our previous work revealed that depleting or chemically inhibiting it significantly suppressed the migration, invasion, and self-renewal properties of triple-negative breast cancer (TNBC) cells in vitro (32). Conversely, increased CPSF73 levels enhanced self-renewal capacity, suggesting that modulating CPSF73 may provide a new strategy for cancer treatment by dysregulating 3’ end processing of critical genes and pathways involved in cancer progression and metastasis. A goal of the current study was to identify these genes.

As reported here, we determined the consequences of CPSF73 inactivation in several different cancer cell types treated with JTE-607, a highly specific, small-molecule inhibitor of CPSF73 activity (34, 35). Importantly, we found that in addition to the previously reported impairment of cancer cell proliferation, it also promotes EMT reversal. Using mRNA 3’ end-focused sequencing to globally profile APA changes, we show that JTE-607 induces 3′ UTR lengthening and suppresses the use of intronic PASs. Our data reveals that 3′ UTR lengthening of the mRNAs of several key genes critical for driving EMT and metastasis is correlated with downregulation of the encoded proteins. We further show that for one of these genes, AKT2, shifting the balance of mRNA isoforms toward the long isoform and the subsequent reduced AKT2 protein expression is sufficient to push the cells away from the mesenchymal state, as evidenced by changes in EMT-related markers and impaired cell invasion. These findings indicate a significant functional connection between APA and the EMT process that can be manipulated to reverse EMT in cancer cells, and raise the possibility that interfering with this APA could offer a new therapeutic strategy with potential efficacy across multiple pathways.

## Methods

### Cell culture, JTE-607 treatments, and generation of CPSF73 knockdown cells

Human cell lines, including MDA-MB-231 (triple-negative breast cancer), HepG2 (hepatocellular carcinoma), MCF7 (breast cancer), and A549 (lung cancer), were obtained from ATCC and cultured in Dulbecco’s modified Eagle’s medium (DMEM)/high glucose supplemented with 10% fetal bovine serum (FBS), and 100 U/mL penicillin, and 100 μg/mL streptomycin. Cells were maintained at 37°C in a 5% CO_2_ humidified incubator. Then, during the exponential growth phase (70-80% confluency), cells were detached using 0.05% trypsin/EDTA and seeded at 30% confluency. The following day, cells were treated with JTE-607 (2.5 μM or 10 μM) (Tocris, 5185) for 72 hours before harvest for western blot analysis or total RNA extraction. The media and JTE-607 were refreshed every other day. All cell lines used in this study were routinely tested for Mycoplasma contamination using a Mycoplasma detection kit (R&D Systems, CUL001B), following the manufacturer’s instructions.

MDA-MB-231 cells with inducible CPSF73 knockdown used in this study were generated as previously described (32). Briefly, human small hairpin RNA (shRNA) sequences targeting CPSF73 (shCPSF73 #1, TTGAATTTCTAATTCTTCCACA; shCPSF73 #3, TAATACATGATCTTCCTACTTC) were cloned into the LT3GEPIR vector. An shRNA targeting luciferase was used as a control. The plasmids of interest, together with packaging and envelope plasmids, were co-transfected into HEK293FT cells using FuGENE® HD (Promega Corp.), and virus-containing supernatants were used to infect the MDA-MB-231 cells.

Lentivirus-infected MDA-MB231 cells were selected and then induced for shRNA expression using puromycin and doxycycline at a final concentration of 2 mg/mL and 1.5 mg/mL, respectively. Knockdown efficiency of the target protein was validated by Western blotting after three days of doxycycline induction.

### Cell proliferation and IC50 assays

The impact of JTE-607 on cancer cell proliferation rate was evaluated using the sulforhodamine B (SRB) assay as previously described (36). Briefly, cells were seeded at a low confluency of 2-3×10^4^ cells per well in 24-well culture plates and cultured at 37°C in 5% CO_2_. After 24 hours, cells were treated with JTE-607 at low (2.5 μM) and high (10 μM) concentrations or left untreated as a control for 1, 2, 3, 4, and 5 days, with media and JTE-607 refreshed every other day. Following treatment, the media was removed, and cells were washed twice with cold phosphate-buffered saline (PBS) and subsequently fixed with 1 mL of ice-cold 10% Trichloroacetic acid (TCA) in each well for at least 1 h at 4°C. TCA was removed, and cells were then rinsed 5-7 times with 1% acetic acid (vol/vol) and air dried. Fixed cells were stained with 500 mL of SRB (0.4% w/v in 1% glacial acetic acid) (Fisher Scientific, 3520-42-1) for 30 minutes at room temperature (RT) with gentle shaking. Cells were quickly rinsed 5-7 times with 1% acetic acid to remove any unbound dye. Plates were air-dried, and bound SRB was solubilized by adding 1 mL/well of 10 mM Tris base solution (pH 10.5). Absorbance was measured at 492 nm using a GloMax microplate reader (Promega, Madison, WI, USA), and the blank value (10 mM Tris alone) was subtracted from each well. Optical density (OD) values were used to generate proliferation curves for each group. All measurements were performed in triplicate.

To determine the IC50 value for JTE-607, cells were seeded at 10% confluency in 24-well plates in triplicate and allowed to adhere overnight at 37°C and 5% CO_2_. Cells were treated with JTE-607 at increasing concentrations (0, 0.5, 1, 2, 4, 8, 16, 32, and 64 μM) for three days. The remaining steps for fixation, washing, and SRB staining were performed as described above for the proliferation assay.

### Protein extraction and Western blotting

Cells were lysed using RIPA lysis buffer (150 mM NaCl, 50 mM Tris-HCl pH 8.0, 1% Nonidet P-40, 0.5% sodium deoxycholate, 0.1% Sodium dodecyl sulfate (SDS), 5 mM EDTA, 1 mM DTT, 10 mM sodium fluoride, 200 μM sodium orthovanadate and 10 mM β-glycerophosphate), supplemented with 1% phenylmethylsulfonyl fluoride (PMSF) and 1× Halt Protease Inhibitor Cocktail (Thermo Fisher Scientific, 78425). Cell lysate was obtained through cycles of incubation on ice and homogenization by vortexing for 20 min, followed by centrifugation at 16,000 × g for 20 min at 4°C. The resulting supernatants were collected, and protein concentrations were determined using the bicinchoninic acid (BCA) assay according to the manufacturer’s instructions (Thermo Fisher Scientific, 23227).

Protein samples were denatured by adding 4X SDS loading buffer (50 mM Tris-HCl pH 6.8, 2% SDS, 10% Glycerol, 0.1% Bromophenol blue, and 2% β-mercaptoethanol) and heating at 95°C for 10 min. Equal amounts of protein (30–50 μg) from each sample were separated using 12% SDS-polyacrylamide gel electrophoresis and transferred onto a pre-wet polyvinylidene fluoride (PVDF) membrane at 110 V for 45-90 min.

Membranes were blocked with either 5% powdered skim milk or EveryBlot Blocking Buffer (Bio-Rad, 12010020) and then incubated with primary antibodies at 4°C overnight. Membranes were washed three times with Tris-base buffer containing 1% TWEEN 20 (TBS-T) and incubated with appropriate horseradish peroxidase (HRP)-conjugated secondary antibodies (1:10,000) for 1 h at RT. Chemiluminescence signals were developed using an enhanced chemiluminescence (ECL) substrate (Thermo Fisher Scientific, 34578) and visualized using the Syngene and LI-COR Odyssey imaging systems. Western blot images were quantified and normalized to glyceraldehyde-3-phosphate dehydrogenase (GAPDH) and alpha-tubulin expression levels as internal reference proteins using ImageJ software. A list of antibodies used in this study is provided in Supplementary Table S1.

### Real-time PCR analysis (RT-qPCR)

Total RNA was extracted using Trizol reagent (Invitrogen, 15596018) and the Direct-zol^TM^ RNA Miniprep Kit (Zymo Research, R2050), followed by DNase I treatment (Zymo Research, E1011A) to eliminate genomic DNA contamination according to the manufacturer’s instructions. RNA quantity and quality were determined using Nanodrop (Thermo Fisher Scientific) and agarose gel electrophoresis. cDNA was synthesized from 1 μg of total RNA from each sample using LunaScript RT SuperMix Kit (New England BioLabs (NEB), E3010L). To perform RT-qPCR, Luna Universal qPCR Master Mix Kit (NEB, M3003L) and specific primers were used and subjected to initial denaturation of 95°C for 60 seconds, followed by 35-40 amplification cycles of 95°C for 15 seconds and 60°C for 30 seconds, conducted on a CFX Opus 96 Real-Time PCR System. Relative transcript levels were normalized to GAPDH as an internal reference gene. Primers targeting the open reading frame (ORF) and sequences just upstream of the distal polyadenylation site (dPAS) were used to detect total and long transcripts, respectively. Relative transcript levels were calculated using the canonical 2−ΔCT method to determine the ratio of long 3′ UTR transcripts to total transcripts. The ratio of long to total transcripts was used to assess 3′ UTR lengthening (increased ratio) or shortening (decreased ratio) in JTE-607-treated cells compared to the controls.

All RNA samples used in this study were routinely tested for Mycoplasma contamination by RT-qPCR using mycoplasma-specific primers. PCR primer sequences used in this study are provided in Supplementary Table S2. Primer specificity was verified by standard PCR reaction using HotStarTaq Plus Master Mix (QIAGEN) according to the manufacturer’s instructions. The amplified products were assessed by electrophoresis on agarose gels to confirm the presence of a single band of the expected size.

### RNA sequencing by QuantSeq

Total RNA was processed for library preparation using the Lexogen 3′ mRNA-Seq QuantSeq FWD kit according to the manufacturer’s instructions and using poly(A)+ selection to enrich for mRNAs. Quality control, library preparation, and sequencing were performed by Admera Health (South Plainfield, NJ, USA). cDNA libraries were sequenced on an Illumina HiSeq machine (2 × 150 nt) at Admera Health.

### Differential gene expression analysis

The QuantSeq FWD data were processed according to the manufacturer’s recommendation. The read 1 data of the samples was first trimmed using the BBtools (37) to remove adapters and polyA sequence, and then mapped to the human hg38 genome using STAR (v2.7.7a) (38). The number of reads mapped to each gene was then quantified using featureCounts (v1.6.3) (39), and only genes with more than five reads per sample were included in further analyses. Read counts were normalized to the total mapped reads to the genome. PseudoCount of 1 was applied to prevent infinity values in the ratio calculation. The significance of expression difference was assessed by DESeq2. All *P* values from DESeq2 were adjusted by the Benjamini-Hochberg (BH) method to control the false discovery rate. BH-adjusted *P* value < 0.05 was considered significant. A fold change of 1 was additionally applied to select regulated genes.

### APA analysis

APA analysis was conducted using a polyA_DB-based pipeline for APA analysis as described in our previous study (40). Briefly, adapter and poly(T) sequences from QuantSeq FWD raw read 2 data were trimmed and aligned to the human (hg38) polyA_DB reference (41) (−100nt, polyA_DB-annotated-PolyA Site (PAS), +25nt) using STAR (v2.7.7a), and only uniquely mapped reads were kept. The last aligned position of each mapped read was compared to polyA_DB-annotated PASs, allowing ± 24 nt flexibility. Matched reads were considered as pA site-supporting (PASS) reads and were used for further APA analysis. UCSC genome browser plots shown in this study were generated based on PASS reads.

For 3′ UTR APA analysis, the two most frequently used PASs in the 3′ UTR were designated as proximal PAS (pPAS) and distal PAS (dPAS) isoforms for comparison. For intronic APA analysis, all intronic isoforms were combined and compared to all isoforms using PASs in the terminal exon (TPA isoforms). Relative Expression Difference (RED) values were calculated as the changes in log2-ratio of isoform abundance (dPAS isoform vs. pPAS isoform) between JTE-607-treated and control samples. Significant APA events were defined as those with RED > log2(1.2) and BH-adjusted *P* value < 0.05 (DEXSeq) (42). Genes were classified as “shortened” or “lengthened” when RED values were ≤ log2(1.2) or ≥ log2(1.2), respectively.

The presence of RBP-binding sites was identified as described in our previous work (40). We overlapped results from RBPmap (43), a tool that predicts RBP-binding based on motif sequences, with CLIP-seq data from the POSTAR3 (44), database to identify RBP-binding sites within 200 bp upstream and downstream of proximal and distal PASs, as well as in the region spanning 175 bp downstream of the proximal site and 175 bp upstream of the distal site. For the ERBB2 gene, the region spanning 200 bp upstream of the proximal PAS and 200 bp downstream of the distal PAS was analyzed for RBP-binding, as the PASs are only 82 bp apart.

### Antisense morpholino oligonucleotide (AMO) treatments

For AKT2 AMO treatment, MDA-MB-231 cells were cultured in DMEM/high glucose supplemented with 10% FBS, 100 U/mL penicillin, and 100 μg/mL streptomycin at a density of 2× 10^6^ per well in 6-well plates for 24 hours. Then, cells (with over 80% confluency) were transfected with AMOs (Gene Tools, LLC) at a concentration of 5 μM/mL using Endoporter (6 μL/mL) (Gene Tools, LLC) according to the manufacturer’s protocol. RNA samples were collected 24 hours post-transfection. As described above, the isolated RNA was then analyzed by RT-qPCR using primers specific to the open reading frame (ORF) of AKT2 mRNA, which represents the total transcripts, and a primer pair targeting sequences just before the dPAS, representing the long transcripts. The ratio of long 3′ UTR transcripts to total transcripts was calculated.

For western blot analysis of AKT2 protein expression and EMT markers, protein samples were harvested 48 hours post-transfection. The following AMO sequences were used:

**Control AMO:** 5’CCTCTTACCTCAGTTACAATTTATA3’

**AKT2_PAS1 AMO:** 5’TGAGTGTCTTTATTGCTTGTACCGT3’

Control AMO is a negative control Morpholino oligo that targets a human beta-globin intron mutation that causes beta-thalassemia.

### Cancer cell invasion transwell assay

To evaluate the invasion potential of MDA-MB-231 cells following AKT2 AMO treatment, Matrigel was thawed at 4°C overnight and diluted with cold serum-free DMEM according to the manufacturer’s instructions. Then, the upper chambers of the 24-well transwell inserts with 8 μm pores were coated with 80 μL of diluted Matrigel and incubated at 37°C overnight. Excess coating buffer was carefully removed from the chambers without disturbing the Matrigel layer covering the membranes.

Following 48 hours of AKT2 AMO treatment, cells were detached using trypsin/EDTA, washed, and suspended at a concentration of 1 × 10^5^ cells per 200 μL medium supplemented with 1% FBS, and seeded on the upper chambers of the Matrigel-coated transwell inserts. The inserts were then placed into lower chambers filled with 600 μL of high-glucose DMEM containing 20% FBS and incubated at 37°C for 72 hours.

To stain invaded cells that transversed the Matrigel and porous membrane, the inserts were removed from the 24-well plate and washed twice with PBS. Non-invaded cells were scraped off from the upper side of the membrane using a cotton swab soaked in warm DMEM. The invading cells were fixed with ice-cold methanol for 15 min at 4°C and then washed twice with PBS and stained with 0.5% crystal violet dye for 15 minutes. After washing with PBS, the stained invading cells were visualized under an inverted microscope. Images of each group were captured; the number of invading cells was quantified using ImageJ software. Data were analyzed using GraphPad Prism 8.0 and presented as the mean of relative cell invasion% ± SEM for four independent experiments. The following formula was used to calculate the relative cell invasion percentage:

Relative Cell Invasion% = (Number of invaded cells/ Mean number of invaded cells in the control group)×100.

### Statistical analysis

All experiments were conducted with three or more independent biological replicates. Statistical analyses were performed using GraphPad Prism Software 8.0 (GraphPad Software Inc., La Jolla, CA, USA). A Student’s *t*-test was applied to evaluate the significance between groups. Data are presented as mean ± SEM. A *P*-value <0.05 was considered significant, with significance levels defined as * = P ≤ 0.05, ** = P ≤ 0.01, and *** = P ≤ 0.001.

## Results

### Inhibition of cancer cell growth by CPSF73 inactivation or depletion correlates with decreased MYC expression and increased p21^WAF1/Cip1^ expression

Analysis of data from the Cancer Genome Atlas (TCGA) shows that the expression of CPSF73 is significantly higher in tumor tissues across many cancers compared to normal tissue (Supplementary Fig. 1A) (45), suggesting a strong dependency of cancer cells on high CPSF73 activity. Clinical prognostic correlation analysis further revealed that this elevated CPSF73 expression is associated with unfavorable outcomes in cancer patients (33, 46, 47). JTE-607 has been shown to strongly suppress the growth of Acute Myeloid Leukemia and Ewing Sarcoma cells by binding to the active site of CPSF73 and blocking its activity (34, 35). Additionally, JTE-607 demonstrated efficacy in suppressing other cancer cell types, including breast, lung, and liver cancers, though with different sensitivity levels to the compound (32, 35, 48–50). To further understand the consequences of CPSF73 inhibition on the growth of cancer cells, we first determined tolerance levels to JTE-607 by measuring the IC50 values of JTE-607 for representative human cancer cell lines, including HepG2 (liver carcinoma cancer), MCF7 (poorly aggressive breast cancer), and A549 (lung adenocarcinoma cancer) cell lines (Supplementary Fig. 1B-D), and used our previous IC50 value of JTE-607 (˃ 50 µM) for MDA-MB-231 (highly invasive Triple Negative Breast Cancer) cells (32). With these cell lines, we could cover a range of cancer behaviors, from less aggressive to highly metastatic, and from different tissue types. Consistent with the findings of Ross, et al. (35), HepG2 cells are very sensitive with an IC50 of 4.2 μM, while the other three were relatively resistant, with IC50 values around 40 μM or greater. Due to the markedly different tolerance levels to JTE-607, cells were then cultured at concentrations of low (2.5 μM) and high (10 μM) doses of JTE-607, and proliferation was measured over five days. JTE-607 significantly attenuated proliferation in all cancer cell lines compared to untreated control cells (Figure 1A-D).

**Figure 1:**
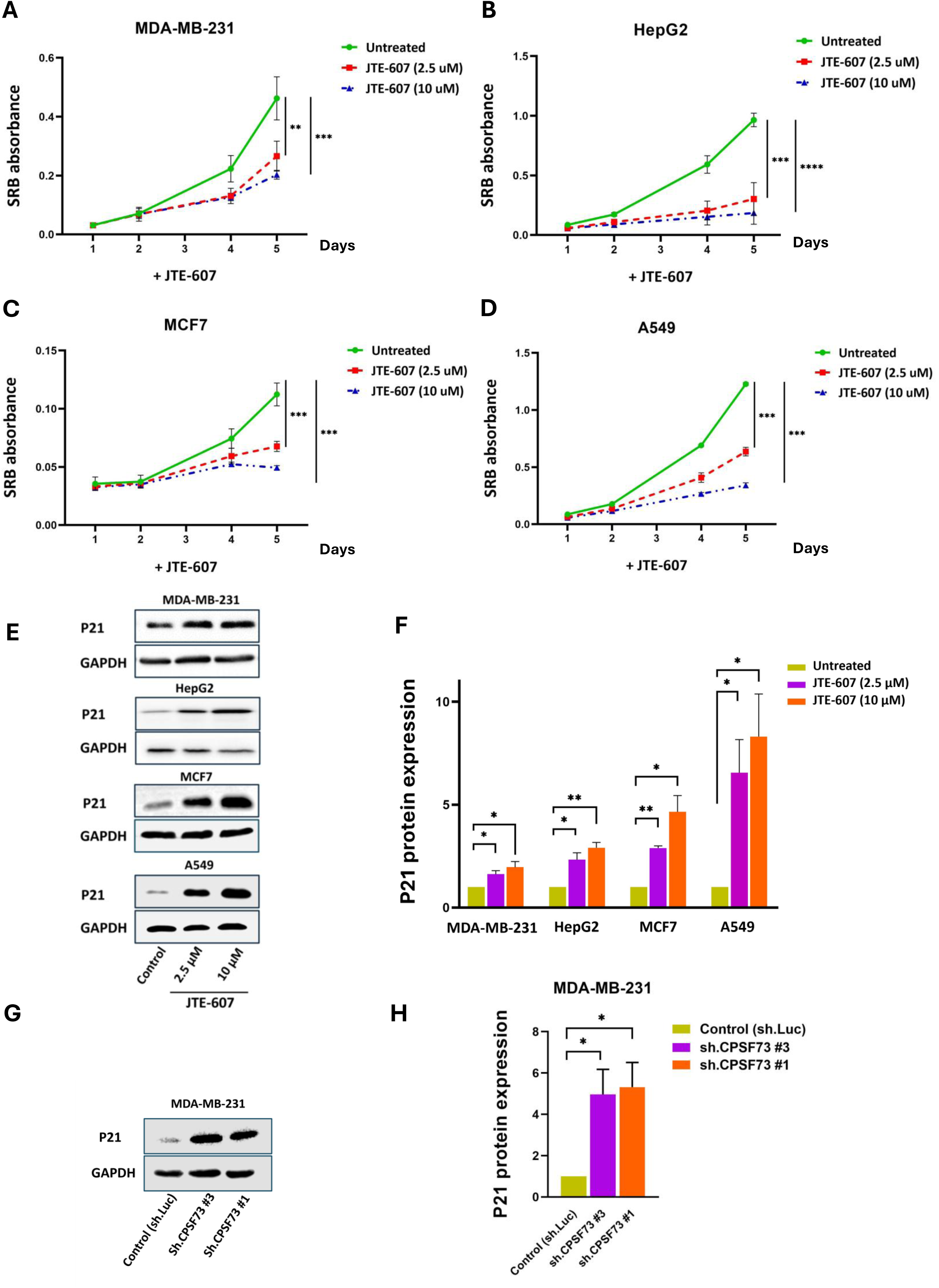
Growth inhibition and increased p21^WAF1/Cip1^ expression following CPSF73 inactivation using JTE-607 in various cancer cell lines. **(A-D)** Cell proliferation curves of cancer cell lines (MDA-MB-231, HepG2, MCF7, and A549) treated with JTE-607 (2.5 and 10 µM, as indicated) compared to untreated control cells over 5 days, as determined by the Sulforhodamine B (SRB) assay, which measures absorbance as optical density (OD) values at 490 nm. **(E)** Representative western blot images and **(F)** corresponding densitometric bar graph show a significant increase in the protein expression levels of p21^WAF1/Cip1^ following CPSF73 inactivation in cancer cells compared to the controls. **(G)** Representative western blot image and **(H)** corresponding densitometric bar graph exhibit a significant increase in p21 protein expression following CPSF73 depletion in MDA-MB-231 cells compared to the control. GAPDH was used as a protein loading control for normalization. Data are presented as mean ± SEM from at least three independent experiments (*p<.05, **p<.01, and ***p<.001).

Checkpoint mechanisms, such as the cyclin-dependent kinase inhibitor p21, also known as CDKN1A or p21^WAF1/Cip1^, promote growth inhibition in response to anti-proliferative stimuli (51). Cancer cells treated with JTE-607 exhibited a significant increase in p21 protein expression compared to their corresponding controls (Figure 1E-F). To further confirm that the increased expression of p21 was triggered by CPSF73 inhibition rather than any possible off-target effects of JTE-607, we used MDA-MB-231 cells with doxycycline-inducible Knockdown (KD) of CPSF73 that were previously generated in our lab (32). CPSF73 depletion in these cells was confirmed by examining the protein level of CPSF73 (Supplementary Fig. 1E). We tested whether CPSF73 KD could also regulate the protein expression of p21 in a similar manner. P21 protein was significantly increased in both CPSF73 KD cells, consistent with our observation in cells with CPSF73 inactivation using JTE-607 (Figure 1G-H).

CPSF73 depletion or inactivation causes defects in mRNA cleavage and leads to global transcriptional readthrough of PASs (32, 35). One of the target genes showing readthrough transcription following CPSF73 downregulation in MDA-MB-231 cells is MYC (33), which regulates p21 expression through direct interaction with transcription factors such as Sp1 and Sp3 near the transcription start site, thereby blocking their ability to activate p21 transcription (52). To assess MYC readthrough transcription following CPSF73 depletion or inactivation in all of our cancer cell lines, we used previously described primers detecting unspliced (US) transcripts and RNA sequence spanning the PAS (Span) of MYC mRNA and monitored MYC transcript levels by RT-qPCR. All cells treated with JTE-607, as well as cells with CPSF73 KD, showed increased levels of RNA spanning the PAS (“Span product”) when normalized to the unspliced *MYC* transcript “US” (Figure 2A-B). This reduction in PAS cleavage likely impairs MYC pre-mRNA maturation, which would lead to decreased MYC protein expression. Indeed, we detected such a decrease in MYC protein in MDA-MB-231 cells following JTE-607 treatment or CPSF73 depletion (Figure 2C-F). Therefore, one possible contributor to the observed increase in p21 expression could be a decrease in MYC protein caused by defective processing at the MYC PAS induced by CPSF73 depletion or inactivation.

**Figure 2:**
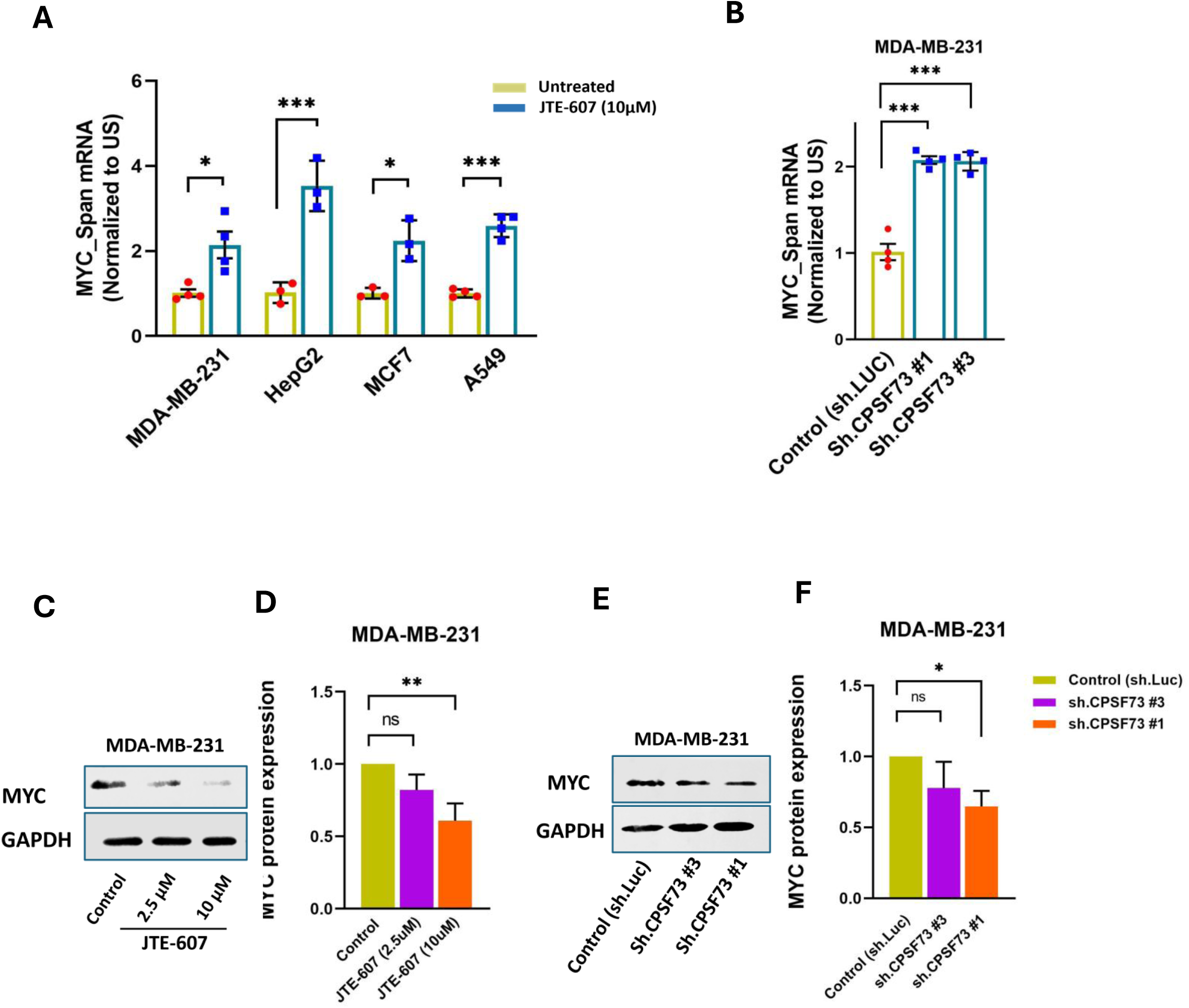
CPSF73-dependent impairment of 3’ end processing and transcriptional readthrough of PASs in MYC. **(A)** Analysis of MYC 3’ end processing in all four cancer cell lines following JTE-607 treatment and **(B)** MDA-MB-231 cells after CPSF73 KD. “Span” values are expressed relative to control cells after normalizing to unspliced (US) RNA levels. **(C and E)** Representative western blot images and **(D and F)** corresponding densitometric bar graphs show a significant decrease in MYC protein expression following CPSF73 inactivation or depletion in MDA-MB-231 cells compared to the control. Data are representative of at least three independent experiments (*p<.05, **p<.01 and ***p<.001).

Together, these results confirm that chemical inhibition of CPSF73 by JTE-607 impairs the proliferation of our chosen cancer cell lines and is accompanied by decreased MYC and increased p21^WAF1/Cip1^ expression.

### CPSF73 inactivation moves cancer cells away from the mesenchymal state and alters gene expression profiles in MDA-MB-231 cells

Cancer cell invasiveness and metastasis rely on EMT, and several studies suggest that alterations in mRNA 3’ end processing can regulate EMT process. Mapping of the alternative polyadenylation landscape has shown extensive APA changes during EMT of normal mammary epithelial cells (53). Our previous findings revealed that depletion or catalytic inhibition of CPSF73 using JTE-607 suppressed migration and significantly reduced invasiveness in MDA-MB-231 cells, a highly metastatic human triple-negative breast cancer cell line (32). Additionally, increased expression of CPSF73 promoted uncontrolled cell proliferation, migration, and metastasis of hepatocellular carcinoma (47).

To further investigate how inhibition of CPSF73, the endonuclease central to the C/P machinery, influences EMT characteristics in cancer cells, we examined the expression of core EMT factors in all four cancer cell lines treated with JTE-607 as well as MDA-MB-231 cells with CPSF73 KD. CPSF73 inactivation pushed the cells away from the mesenchymal state, as evidenced by an increase in the expression of the E-cadherin epithelial marker that was especially strong in MDA-MB-231 and HepG2 cells, along with a significant decrease in the expression of the Zeb1 and Twist2 EMT-driver transcription factors (Figure 3A-H). The N-cadherin mesenchymal marker decreased in all cell lines except A549, and Vimentin, another mesenchymal marker, decreased in HepG2 and MCF-7 cells, but not in MDA-MB-231 and A549 cells. CPSF73 KD cells also exhibited a similar trend, showing increased E-cadherin expression and decreased N-cadherin and Zeb1 expression, whereas Twist2 and Vimentin were not significantly affected (Supplementary Fig. 2A-B). The EMT reversal was somewhat less effective in CPSF73 KD cells than in those treated with JTE-607, probably because the remaining CPSF73 expression after KD could still compensate for its function. In summary, while there were differences likely due to the origin and mutational background of the different cell lines, all cell lines showed significant changes in three or more EMT markers.

**Figure 3:**
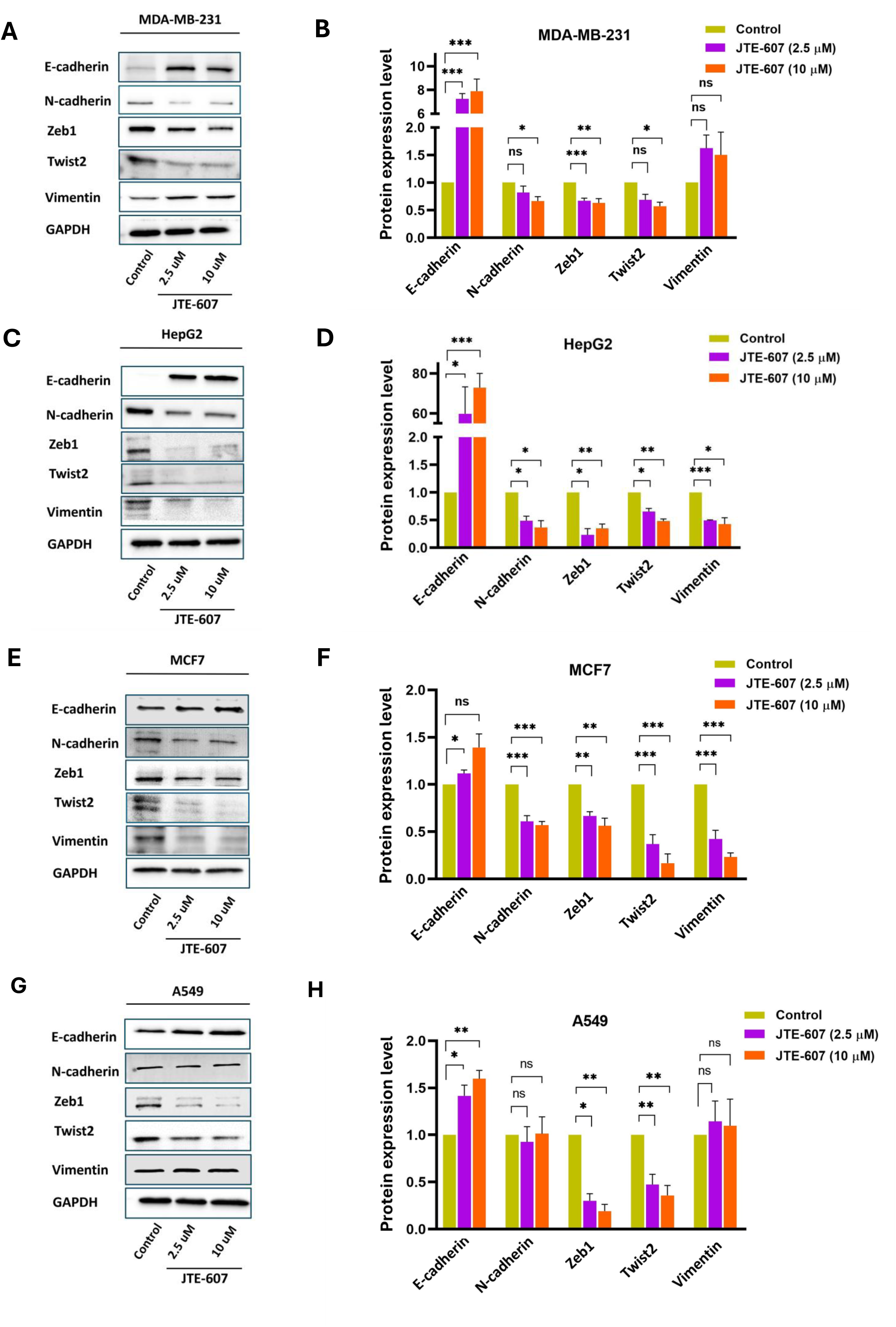
Western blot analysis demonstrating altered expressions of EMT-related proteins in cancer cell lines following JTE-607 treatment. Representative western blot images and corresponding densitometric bar graphs quantifying the expression of EMT-related markers in MDA-MB-231 **(A** and **B)**, HepG2 **(C** and **D)**, MCF7 **(E** and **F)**, and A549 **(G** and **H)** cancer cells treated with JTE-607 for three days, compared to untreated controls. GAPDH was used to control the protein loading and normalize samples. Data are presented as mean ± SEM from at least three independent experiments (*p<.05, **p<.01, and ***p<.001).

Based on these findings, we hypothesized that changes in mRNA 3’ end processing are likely involved in the suppression of EMT driven by JTE-607 treatment or CPSF73 depletion. To test this hypothesis and to elucidate the consequence of the cleavage defect on the global APA landscape and gene expression, we conducted mRNA 3’ end-focused sequencing (QUANT-Seq) on MDA-MB-231 cells with or without JTE-607 treatment (10 µM for 72 hours). We chose the highly aggressive MDA-MB-231 TNBC cell line for this part of the study to build on our previous work elucidating the role of CPSF73 in the invasion and self-renewal capabilities of these cells (32). This global sequencing analysis enables the simultaneous determination of both gene expression and APA changes (54, 55). To compare the expression profiles of JTE-607-treated cells with the control, we performed differential gene expression (DGE) analysis. A scatter plot of the distribution of differentially expressed genes revealed widespread gene expression changes following JTE-607 treatment (Figure 4A). Notably, the number of significantly upregulated (2,654), and downregulated (2,583) genes was nearly equal. The complete list of DEGs is available in Supplementary File 1.

**Figure 4:**
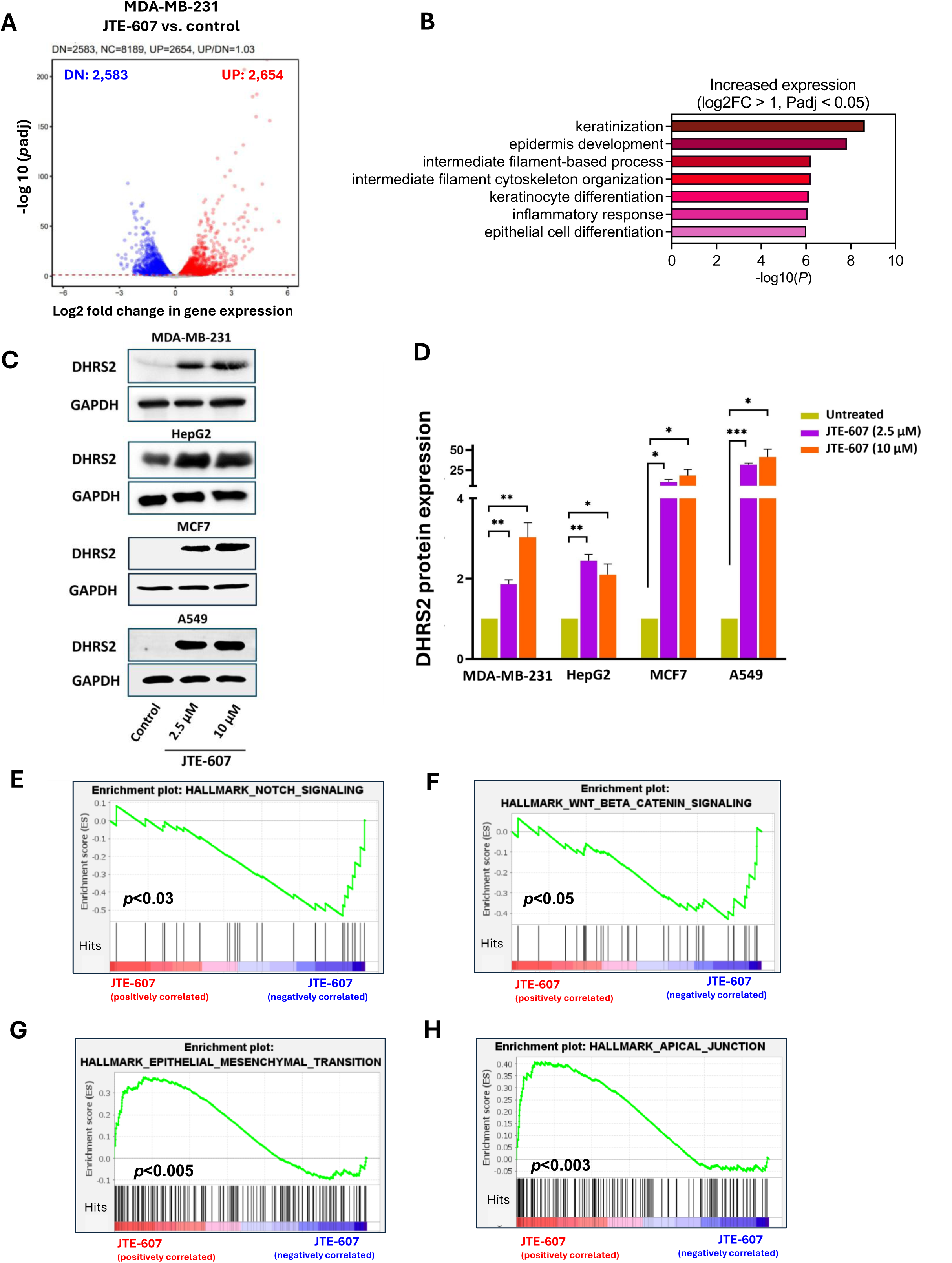
RNA sequencing representing differentially expressed genes and gene set enrichment analysis (GSEA) following CPSF73 inhibition using JTE-607 in MDA-MB-231 cells. **(A)** Scatter plot illustrating the distribution of differentially expressed genes (DEG) using DESeq2 in JTE-607-treated MDA-MB-231 cells versus untreated controls, with red and blue dots representing upregulated (UP) and downregulated (DN) genes, respectively that have a DGE regulation cutoff of log2-fold change ≥ 1 or ≤-1, and p-adjusted value (padj) < 0.05), and gray dots representing unchanged genes. **(B)** Gene Ontology analysis of genes with increased expression after JTE-607 treatment of MDA-MB-231 cells. The top 7 GO terms are presented. **(C)** Representative western blot images and **(D)** corresponding densitometric bar graphs show a significant increase in the protein expression levels of DHRS2 following CPSF73 inactivation in cancer cells compared to the controls. GAPDH was used as a protein loading control for normalization. Data are presented as mean ± SEM from at least three independent experiments (*p<.05, **p<.01, and ***p<.001). **(E, F, G, and H)** GSEA Hallmark analysis revealed enriched gene sets. The GSEA-enrichment plots show negatively enriched Notch **(E)** and WNT **(F)** signaling pathways and positively enriched EMT**(G)** and Apical Junction (**H)** hallmark gene sets in the JTE-607-treated group compared to the controls. Genes are ranked by expression fold change.

In agreement with our western blot observations of changes in EMT-related proteins, gene ontology (GO) analysis of upregulated genes in MDA-MB-231 cells treated with JTE-607 showed that the most prominent GO categories associated with increased gene expression were related to epithelial differentiation (Figure 4B). Consistent with our proliferation and EMT markers analysis, DHRS2, a protein involved in metabolic regulation, inhibition of cancer cell growth, and suppression of metastasis (56, 57) was the most upregulated gene in JTE-607-treated MDA-MB231 cells. Western blot analysis further confirmed a marked increase in DHRS2 protein levels in treated cells compared to their corresponding controls (Figure 4C and D).

To further understand the impact of CPSF73 inactivation, we conducted Gene Set Enrichment Analysis (GSEA) (58) to identify the most significant overrepresented or modulated canonical biological pathways, and to determine whether gene sets related to invasion and EMT features of cancer cells are systematically altered following JTE-607 treatment. For this analysis, all genes were ranked by log2-fold change according to the extent of their differential expression in the RNAseq data, and Hallmark gene sets derived from the Molecular Signatures Database (MSigDB) were used. The most affected pathways are presented in Supplementary File 2. Notably, the Notch and WNT signaling pathways, which promote EMT (59, 60), were negatively enriched (down-regulated) among the overrepresented gene sets in JTE-607-treated cells compared to control (Figure 4E and F). Conversely, at the other extreme, the EMT hallmark gene set, which contains genes that consistently showed coordinated up- or down-regulation with the EMT phenotype across multiple expression datasets, and the Apical junction (a structure mediating contact between adjacent epithelial cells) gene sets were positively enriched in JTE-607-treated cells (Figure 4G and H), supporting the hypothesis that CPSF73 inactivation promotes the reversion of EMT.

### JTE-607 primarily induces 3′ UTR lengthening and suppresses the use of intronic PASs

Shifting the position of the PAS through APA in the last exon results in mRNA isoforms with varying 3′ UTR lengths, while APA in intronic sites produces transcripts with different coding sequences. These changes can consequently affect mRNA stability, translation, localization, and protein function (14, 18, 61, 62). To identify differential usage of PASs, we assessed APA regulation by calculating Relative Expression Difference (RED) values, representing the usage of the proximal PAS relative to the distal PAS, as described in the Methods section. Mapping the PAS usage from QUANT-Seq data showed APA changes of 943 genes with lengthened 3′ UTRs, 398 genes with shortened 3′ UTRs, and 3121 genes with unchanged 3′ UTR APA, resulting in a lengthening to shortening ratio of 2.4 (Figure 5A).

**Figure 5:**
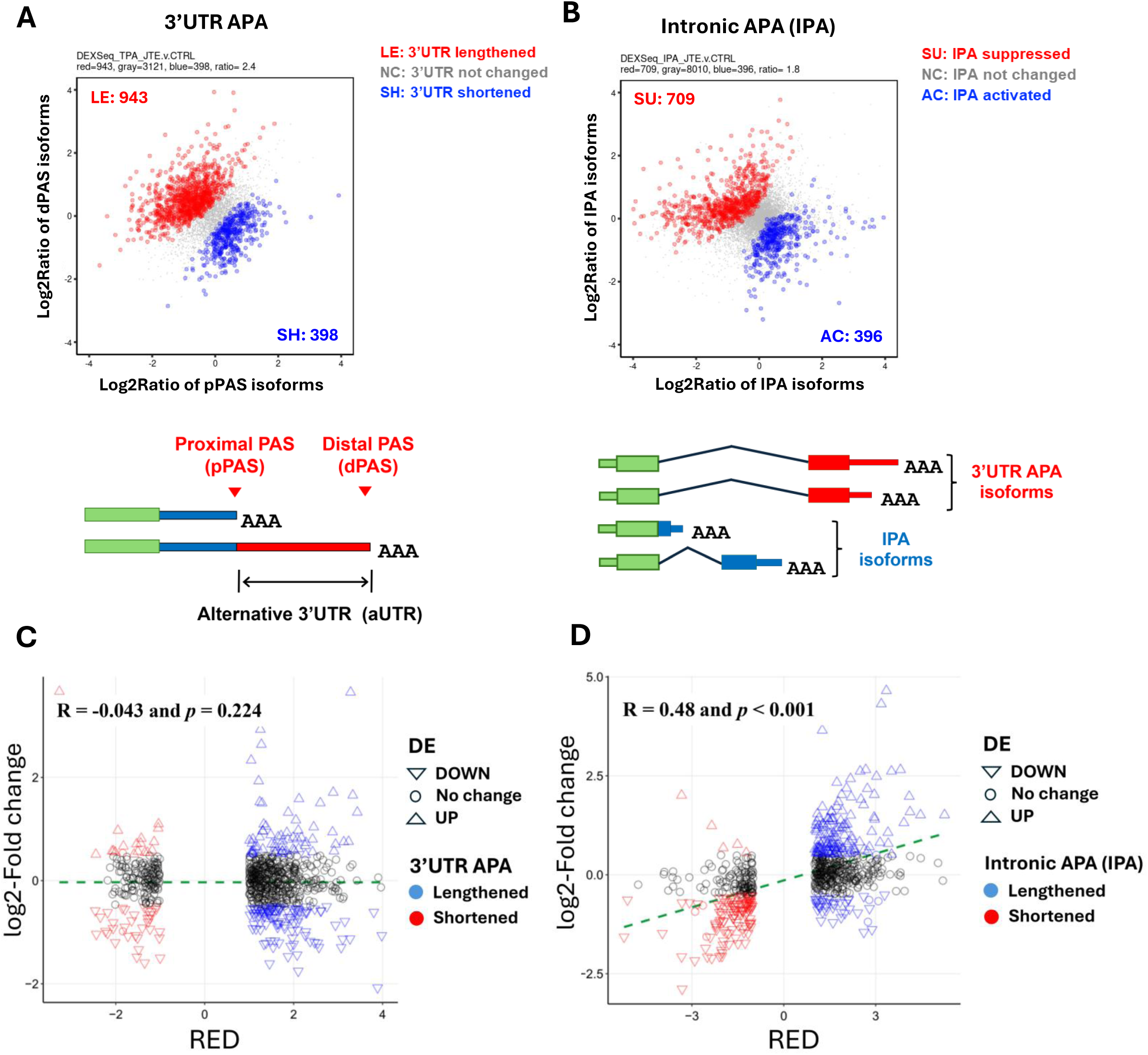
Analysis of the global APA landscape reveals that JTE-607 treatment primarily causes 3′ UTR lengthening and intronic PAS suppression. mRNA 3’ end-focused sequencing was performed on RNA samples obtained from MDA-MB-231 cells treated with 10 µM JTE-607 for 72 hours compared to untreated controls. **(A** and **B)** Scatter plots showing significant and widespread differences in 3′ UTR and intronic APA between JTE-607-treated and control MDA-MB-231 cells. dPAS, distal pA site; pPAS, proximal pA site. **(C** and **D)** Scatter plots show correlation analysis between RED and Log2-Fold expression changes for genes exhibiting 3′ UTR lengthening or shortening **(C)**, and intronic polyadenylation (IPA) changes **(D)**. Upward-pointing triangles represent upregulated genes (*p*-adjusted < 0.05, log2-fold change > 0.5), while downward-pointing triangles indicate downregulated genes (*p*-adjusted < 0.05, log2-fold change < −0.5). Black circles represent unchanged genes. Triangle outline colors indicate whether genes were shortened (red) or lengthened (blue) in their 3′ UTRs or IPA usage. The correlation between log2-fold changes and RED values is visualized by a green dashed line of best fit, with Spearman’s correlation value and corresponding p-value shown in the top left of each graph.

Analysis of intronic APA (IPA) changes after JTE-607 treatment revealed decreased intronic PAS usage, or suppression, in 709 target genes, and increased intronic PAS usage, or activation, in 396 target genes, with a suppression to activation ratio of 1.8, and 8010 genes with no changes in intronic PAS usage (Figure 5B). These data highlight significant and widespread changes in both 3′ UTR and intronic APA events upon inhibiting the mRNA cleavage activity of CPSF73 in MDA-MB-231 cells. Detailed output for all APA events is provided in Supplementary Files 3 and 4.

To examine the potential consequences of APA changes on differential gene expression changes of affected genes, the RED values of genes exhibiting shortening or lengthening due to APA or IPA changes were plotted against their log2-fold expression changes. We found that genes undergoing 3′ UTR APA did not show a significant correlation with gene expression (R = −0.043 and *p* = 0.224) (Figure 5C). However, genes with intronic APA changes showed a modest but statistically significant correlation with expression (R = 0.48 and *p* ˂ 0.001). This suggests an association between IPA activation and decreased expression, as well as IPA suppression and increased expression (Figure 5D).

To determine how APA changes contribute to the EMT reversal, EMT-related genes were extracted from the BioCarta, MSigDB, and GeneCard databases (Supplementary File 5). An overlap analysis of EMT-related gene set with the set of genes with 3′ UTR APA changes, defined by RED value ≤ −1 or ≥ 1, following JTE-607 treatment identified 42 genes in common (Supplementary Fig. 3A). Based on the magnitude of the 3’ end processing changes after JTE-607 treatment and well-known functions in EMT, a subset of genes was selected for further validation by qPCR and western blot analysis. Representative UCSC genome browser plots of RNA sequencing tracks for lengthened genes after JTE-607 treatment in MDA-MB-231 cells are shown in Figure 6A, C, E, and G for TGFBR1, SMAD4, TBX2, and PAK1, with additional examples (CD44 and ERBB2) shown in Supplementary Fig. 3B and D. For each of these genes, the change in reads at the annotated PASs is consistent with the APA shifts predicted by gene’s RED values.

**Figure 6:**
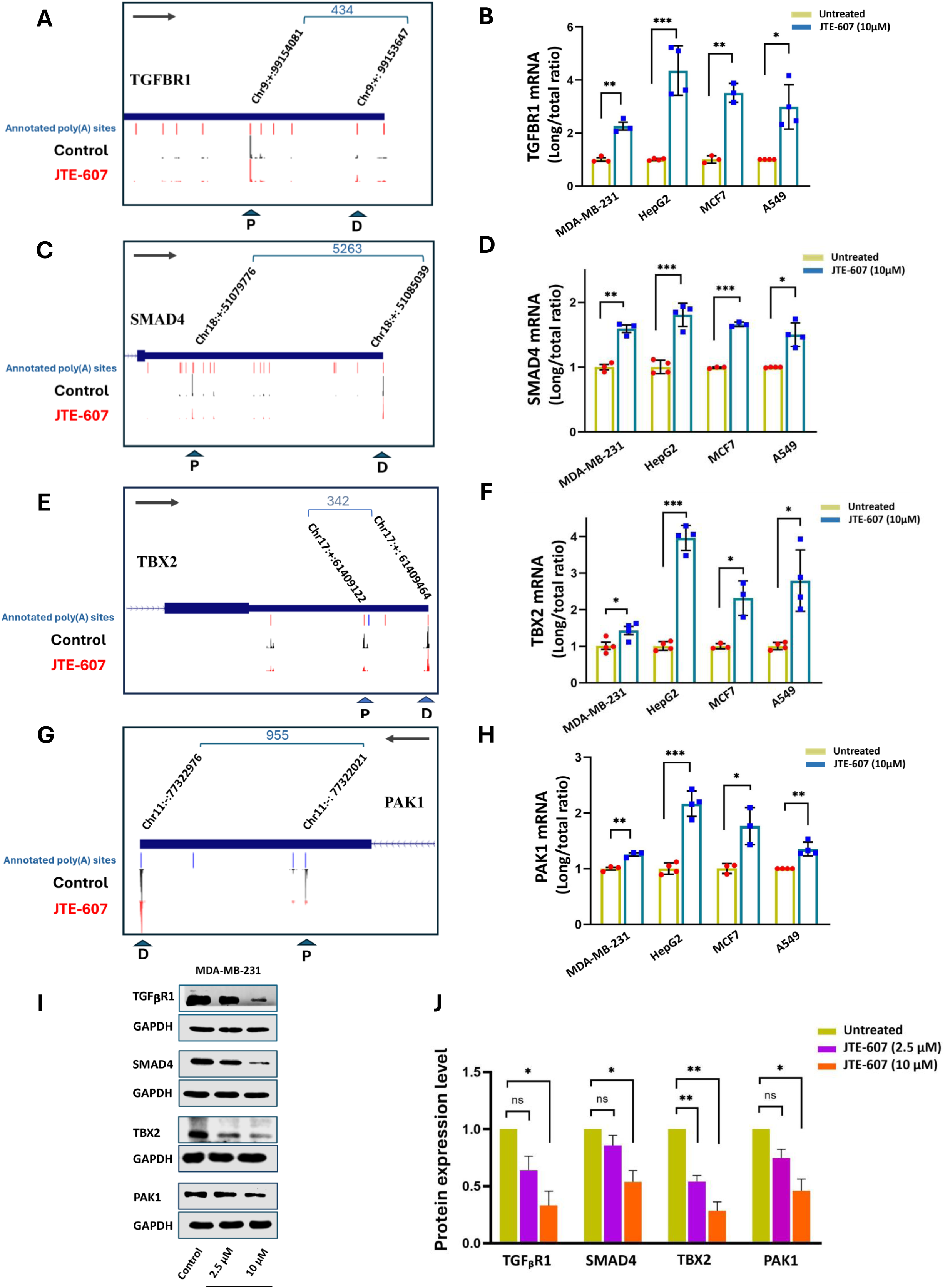
UCSC Genome Browser plots and RT-qPCR validation of EMT-related genes with lengthened 3′ UTRs following JTE-607 treatment. **(A, C, E, G)** UCSC Genome Browser plots of RNA sequencing tracks showing changes in PAS usage, resulting in 3′ UTR profile alterations for lengthened genes TGFBR1, SMAD4, TBX2, and PAK1 in MDA-MB-231 cells. Chromosome co-ordinates and distance between the differentially used PASs are indicated at the top of the plot. The positions of the distal (D) and proximal (P) PASs are indicated with arrows. The arrow inside the box shows the gene direction. **(B, D, F, H)** Corresponding bar graphs showing RT-qPCR validation of 3′ UTR lengthening in all four cancer cell lines after CPSF73 inactivation with JTE-607. Values represent the ratio of long mRNA isoforms to total mRNA (long/total) for untreated (yellow) and treated (blue) samples. Gene expression levels were normalized to GAPDH. **(I** and **J)** Representative western blot images and corresponding densitometric bar graphs show a significant decrease in the protein expression levels of TGFBR1, SMAD4, TBX2, and PAK1 following JTE-607 treatment. GAPDH was used as a protein loading control for normalization. Data are presented as mean ± SEM from at least three independent experiments (*p<.05, **p<.01 and ***p<.001).

To validate the switch between mRNA isoforms resulting from differential usage of cleavage sites observed in QUANT-Seq, we performed RT-qPCR on the lengthened genes using the same RNA samples as those used for sequencing, as well as RNA from HepG2, MCF7, and A549 cells treated with 10 μM of JTE-607. For each gene, we designed a primer set to amplify a common region shared by both isoforms to quantify total mRNA expression, and another primer pair specific to the longer 3′ UTR to detect the long isoform. RT-qPCR analysis indicated that the inhibition of CPSF73 by JTE-607 increased the long/total ratio, indicating 3′ UTR lengthening in these genes in all four cell lines, and supporting the data from the QUANT-seq dataset (Figure 6B, D, F, H, and Supplementary Fig. 3C, E).

Given that 3′ UTR lengthening can introduce additional binding sites for RNA-binding proteins and miRNAs that potentially alter mRNA translation, we were particularly interested in the potential consequences of these APA changes on protein expression levels. To investigate this, we performed western blot analysis on protein samples from MDA-MB-231 cells treated with 2.5 μM and 10 μM of JTE-607 compared to the control. Our results revealed a significant decrease in the protein expression of TGFBR1, SMAD4, TBX2, and PAK1 following JTE-607 treatment, which may be a result of 3′ UTR lengthening in these genes (Figure 6I and J). Together, these findings highlight a strong link between APA events and the reversal of EMT, suggesting that APA may play a critical role in modulating the expression and protein output of key EMT-related genes.

### 3′ UTR lengthening of AKT2 mRNA impairs invasion of MDA-MB-231 cells through downregulation of AKT2 and EMT-related proteins

To probe more deeply into the mechanism underlying EMT reversal through APA, we sought to identify a candidate gene among EMT-related genes exhibiting 3’ UTR APA changes and whose relative expression is altered in the course of CPSF73 inactivation following JTE-607 treatment. According to these criteria, twelve EMT genes were up-regulated and lengthened, none were upregulated and shortened, only one (IDH1) was down-regulated and shortened (Supplementary Fig. 4A-C), and ten EMT genes exhibited both 3’ UTR lengthening and decreased mRNA expression (Figure 7A). The remaining 19 EMT genes with 3’ UTR changes did not show changes in expression, in agreement with the fact that some 3’ UTR changes affect translation or mRNA localization without affecting expression (12–14, 63). Because changes in 3’ UTR length can affect mRNA stability, our analysis focused on the 10 genes found in the intersection of EMT-related genes with those exhibiting both 3’ UTR lengthening and decreased mRNA expression. We wanted to see if manipulating the APA of one of the affected genes would affect protein level, as well as have a functional consequence relevant to EMT. As described below, we selected AKT2 because the sequence around its proximal PAS was sufficiently G/C rich to allow design of a blocking anti-sense oligonucleotide.

**Figure 7:**
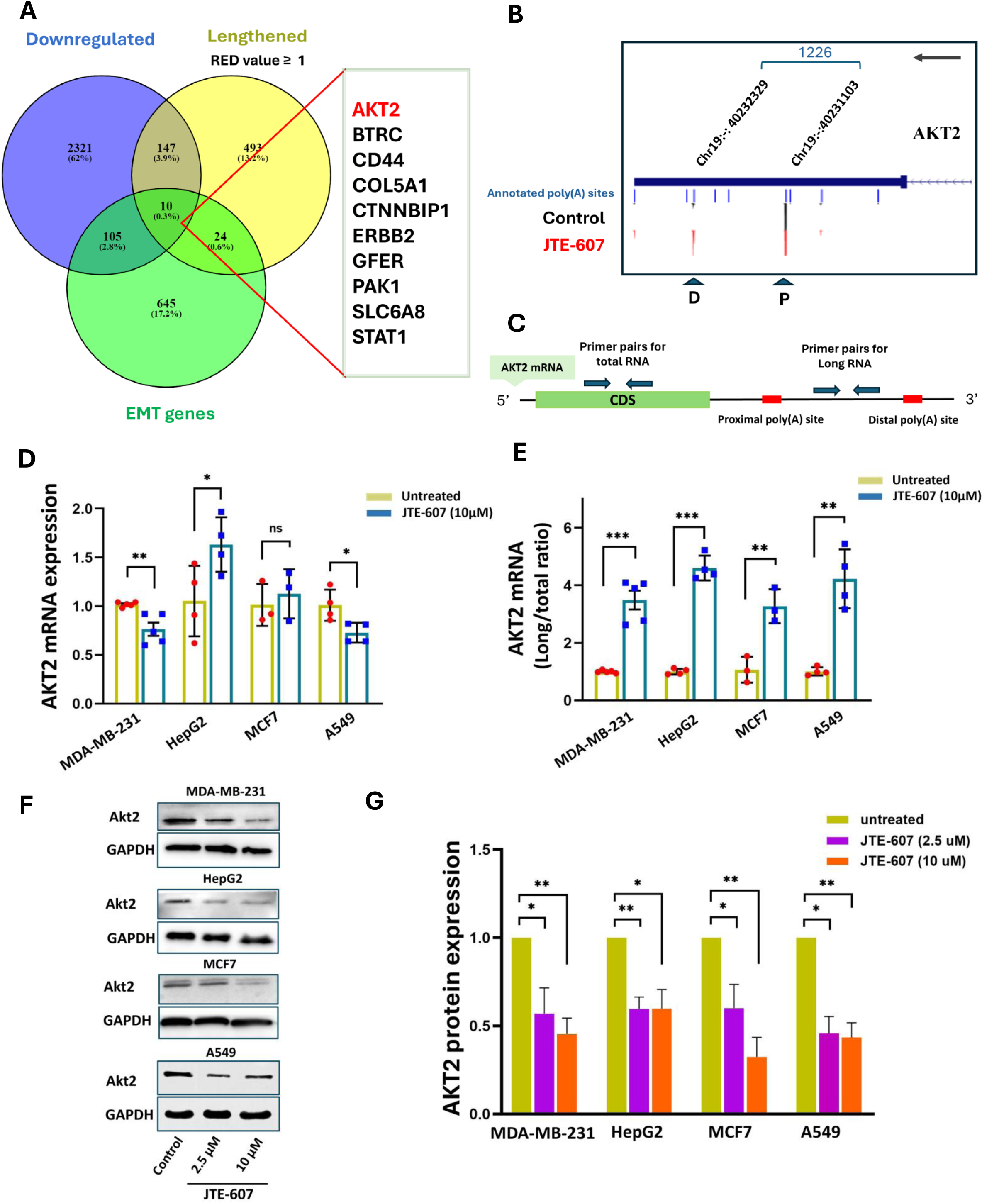
Analysis of AKT2 3′ UTR lengthening and expression following CPSF73 inactivation. **(A)** The Venn diagram shows the intersection of EMT-related genes with genes exhibiting both 3’ UTR lengthening and decreased mRNA expression. **(B)** UCSC Genome Browser plot showing the change in use of the distal (D) and proximal (P) PASs (indicated with arrows) of AKT2 mRNA. Chromosome co-ordinates and distance between the differentially used PASs are indicated at the top of the plot. The arrow inside the box shows the gene direction. **(C)** Diagram of primer binding sites used to confirm 3’ UTR lengthening of AKT2 mRNA by RT-qPCR. **(D** and **E)** RT-qPCR analysis of total mRNA level and changes in the long/total ratio of AKT2 following JTE-607 treatment. Gene expression levels were normalized to GAPDH. **(F** and **G)** Representative western blot images and corresponding densitometric bar graphs showing a significant decrease in the protein levels of Akt2 in cancer cell lines following JTE-607 treatment. GAPDH was used as a protein loading control for normalization. Data are presented as mean ± SEM from at least three independent experiments (*p<.05, **p<.01, and ***p<.001).

AKT2 is an excellent candidate because it is a serine/threonine protein kinase with a well-documented role in promoting both EMT and metastasis (64). AKT2 contributes to migration and invasion and promotes metastasis in various cancer types by regulating multiple functions such as cell adhesion, cytoskeleton rearrangement during migration, matrix metalloproteinases (MMPs) expression, and EMT transcription factors such as Snail (65–71). Silencing of AKT2 through siRNA-mediated depletion or knockout significantly disrupts metastasis and tumor cell growth both in vivo and in vitro (65–67, 69, 72). Additionally, AKT2 expression is elevated in tumor tissue samples compared to normal tissue (Supplementary Fig. 5A). Kaplan–Meier survival analysis further revealed that breast cancer patients with high AKT2 expression have shorter overall survival compared to those with low expression (hazard ratio = 1.14, p < 0.013) (Supplementary Fig. 5B). These studies support a role for AKT2 in cancer progression and invasion, making it a good choice for dissecting the relevance of its APA.

The UCSC genome browser plot of RNA sequencing tracks demonstrated strong 3′ UTR lengthening of AKT2 mRNA, with a RED value of 2.47, after JTE-607 treatment (Figure 7B). To confirm this shift, we detected the usage of the different PASs using one primer pair targeted to the coding sequence to represent the total transcripts, and another pair targeted to the region between the proximal and distal PASs to represent the long isoform (Figure 7C). RT-qPCR analysis showed a significant decrease in total AKT2 mRNA expression in MDA-MB-231 and A549 cells, no change in MCF7 cells, and an increase in HepG2 cells (Figure 7D). Importantly, the 3’ UTR of AKT2 mRNA was lengthened with JTE-607 treatment, as indicated by an increased long/total (L/T) ratio of AKT2 mRNA in all cancer cell lines (Figure 7E). AKT2 mRNA also showed slight lengthening in MDA-MB-231 cells with CPSF73 KD compared to their counterpart control (Supplementary Fig. 5C).

We also performed western blot analysis on protein samples from MDA-MB-231, HepG2, MCF7, and A549 cells treated with 2.5 μM and 10 μM of JTE-607, as well as MDA-MB-231 cells with CPSF73 KD compared to the controls. Our results revealed a significant decrease in Akt2 protein expression in all cell lines following JTE-607 treatment or CPSF73 KD, which may be a result of the 3′ UTR lengthening observed in this gene (Figure 7F-G and Supplementary Fig. 5D and E).

We next investigated whether switching the endogenous AKT2 mRNA from the shorter size to a longer isoform, as observed after JTE-607 treatment, could reverse EMT and reduce the invasion potential of MDA-MB-231 cells. To mimic the observed lengthening of AKT2 mRNAs after JTE-607 treatment, we designed an antisense morpholino oligonucleotide (AMO) complementary to the proximal PAS of AKT2 mRNA to block the usage of this PAS in the endogenous AKT2 gene (Figure 8A). This approach prevents recognition of the proximal PAS by the processing complex, thereby forcing the selection of a more distal PAS and increasing the production of the long isoforms. Treatment with the AKT2 AMO led to a significant decrease in total AKT2 mRNA expression and a marked increase in the relative abundance of the long AKT2 mRNA isoform, as indicated by the increased long to total ratio in MDA-MB-231 cells compared to treatment with a control AMO (Figure 8B and C). This increase in the long/total ratio of AKT2 mRNA was accompanied by a decrease in Akt2 protein expression (Figure 8D and E). While there was no change in E-cadherin, EMT-related markers Zeb1, N-cadherin, and Snail were reduced following AMO treatment. This result aligns with a previous report showing that decreased expression of Akt2 induced by small interfering RNAs attenuates the Snail1 expression induced by TGF-β1 in renal tubular epithelial cells (71).

**Figure 8:**
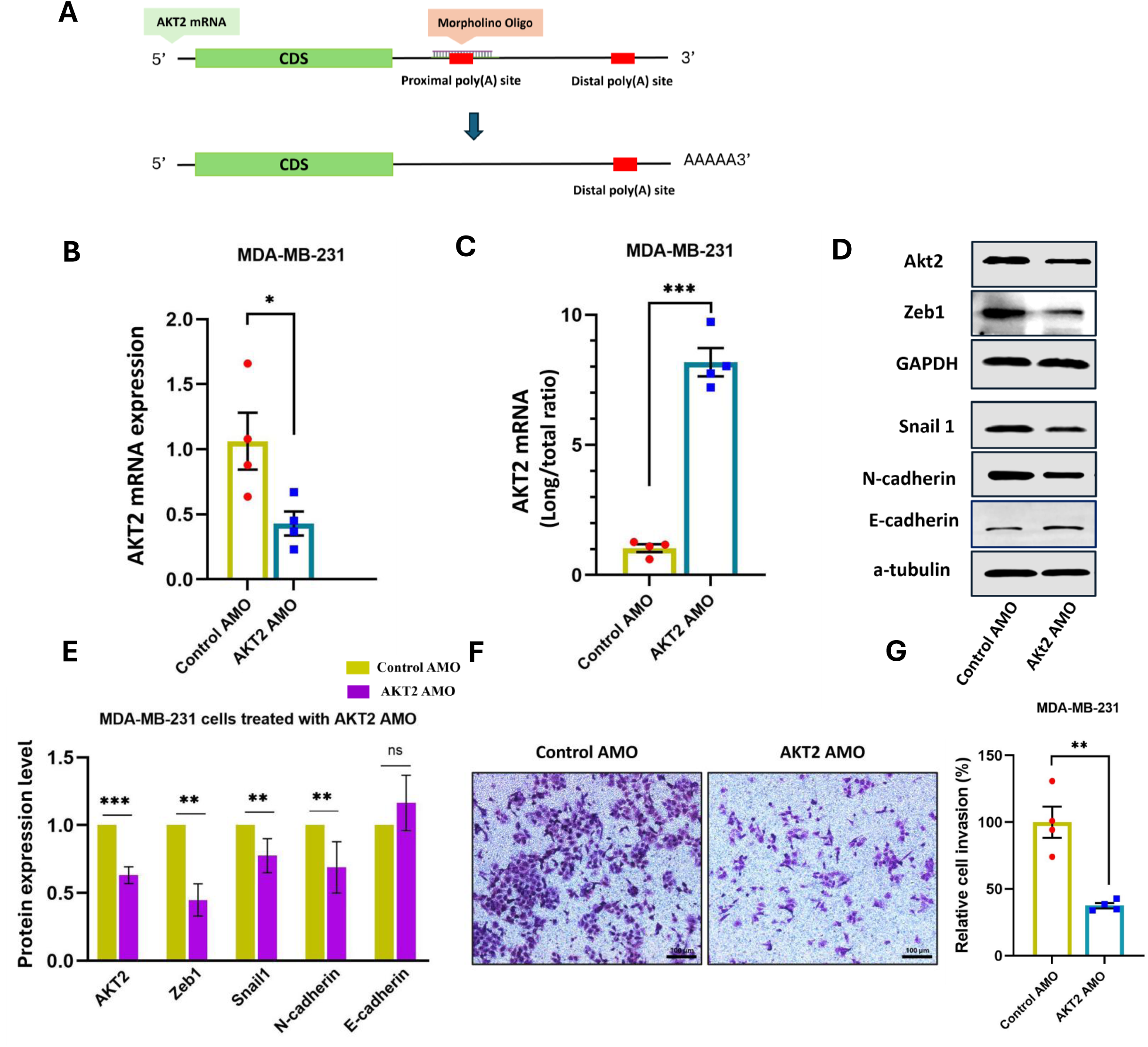
Antisense Morpholino Oligonucleotide (AMO)-induced 3′ UTR lengthening of AKT2 mRNA reverses EMT and reduces invasion in MDA-MB-231 cells. **(A)** The position of the antisense morpholino oligonucleotide (AMO) designed to block the proximal PAS of AKT2 mRNA and shift the endogenous AKT2 mRNA to the longer isoform. **(B** and **C)** RT-qPCR analysis showing total mRNA level, and changes in the long/total ratio of AKT2 mRNA following AMO treatment. Gene expression levels were normalized to GAPDH. **(D** and **E)** Representative Western blot analysis and corresponding densitometric bar graphs quantifying the expression of Akt2 and EMT-related markers following AMO treatment. GAPDH and α-tubulin were used as protein loading controls for normalization. **(F** and **G)** Representative images of the transwell invasion assay and a bar graph depicting the relative cell invasion after JTE-607 treatment, with untreated cells set at 100%. Data are presented as mean ± SEM from at least three independent experiments (*p<.05, **p<.01, and ***p<.001).

To evaluate the functional consequence of AMO-mediated lengthening of AKT2 mRNA, we employed a transwell invasion assay, which monitors the migration of cells through a gel matrix and across a porous membrane. The lengthening of AKT2 mRNA after AMO treatment resulted in a more than 50% reduction in the invasive capacity of MDA-MB-231 cells (Figure 8F and G). These findings establish 3’ UTR length control as a key regulatory mechanism influencing the amount of Akt2 protein available to cancer cells.

## Discussion

Alternative polyadenylation profoundly expands transcriptome diversity by generating distinct transcript isoforms, thereby maintaining diverse physiological processes. Dysregulation of mRNA 3’ end processing is increasingly recognized as a mechanism contributing to carcinogenesis and cancer progression, often through activating proto-oncogenes and the repression of tumor suppressors (21, 24). While recent studies have linked CPSF73 inhibition to slowing cancer cell growth (35, 48, 50), its contribution to regulating EMT characteristics in tumor cells through APA events has remained unexplored. In this study, we report for the first time that APA alterations driven by CPSF73 inactivation can reverse aspects of EMT in cancer cells at a molecular and functional level.

In agreement with other studies (32, 34, 35, 48–50, 73), our work shows that catalytic inhibition of CPSF73 by JTE-607 slows the proliferation of several cancer cell lines with different mutational backgrounds and origins. Consistent with the growth inhibition, we observed increased expression of p21, a key gene inhibitor of cell cycle progression (51, 57), following JTE-607 treatment in all of the cancer cells we tested, as well as in MDA-MB-231 cells depleted of CPSF73. In addition, CPSF73 depletion or inactivation caused increased transcriptional readthrough of the PAS of MYC, which is a well-known negative regulator of p21. This transcriptional readthrough and the decrease in MYC protein level may be one of the drivers leading to p21 upregulation. In addition, PAK1 can directly or indirectly regulate several cyclin-related kinases, Aurora kinases, and checkpoint kinases, and plays a vital role in promoting cell cycle (74). Importantly, PAK1 silencing has been shown to induce p21 expression (75). Therefore, the PAK1 mRNA lengthening and associated reduction in PAK1 protein that we observed following CPSF73 inhibition may also contribute to elevated p21 levels and promote growth arrest. JTE-607 has also been shown to induce S-phase arrest in pancreatic ductal adenocarcinoma (PDAC) cells via impaired histone mRNA processing (48). We propose that a decrease in histones and the suppression of MYC and PAK1 both contribute to p21-mediated growth inhibition in the cancer cell types that we tested.

Importantly, our study points to another node at which CPSF73 inactivation suppresses the cancer phenotype. Consistent with our western blot data showing changes in the expression of EMT-related markers following JTE-607 treatment or CPSF73 depletion, GSEA analysis of our sequencing data revealed that JTE-607 significantly altered the expression of several EMT hallmark genes, resulting in up-regulation of genes related to epithelial differentiation and the apical junction, and in particular, down-regulation of the EMT-promoting Notch and WNT signaling pathways. These signaling pathways have been implicated in the acquisition of EMT and metastasis through crosstalk with several EMT-relevant transcription and growth factors (59, 60). Notch signaling promotes EMT by transcriptional activation of NF-κB, which in turn activates the expression of target genes like Snail1(76). Similarly, Wnt/β-catenin signaling can promote EMT by transcriptional reprogramming of EMT-TFs through a variety of mechanisms, like crosstalk with TGFβ (59). Additionally, both Wnt and Notch signaling stabilize β-catenin expression, further facilitating the upregulation of EMT regulators such as Vimentin, Slug, and Snail1. Conversely, inhibiting these pathways can reverse the EMT process (59, 60); our findings show that CPSF73 inhibition can have a similar effect on EMT.

Emerging evidence suggests that changes in APA profiles may actively drive EMT. For instance, the sustained and elevated expression of ZEB1, a core EMT-TF, is promoted through APA-mediated shortening of its 3’ UTR under chemotherapeutic-induced genotoxic stress (26). Analysis of tumor tissue samples from patients with metastatic non-small cell lung cancer (NSCLC) and NSCLC cell lines revealed a trend toward shorter 3’ UTRs in cancer cells with a more mesenchymal phenotype compared to those with more epithelial features (77). Additionally, widespread 3’ UTR lengthening and shortening have been observed during TGFβ-induced EMT in normal mammary epithelial cells (30). A previous study by our group demonstrated that depletion or catalytic inhibition of CPSF73 reduced migration and invasion in both MDA-MB-468 (Basal A type TNBC) and MDA-MB-231 (Basal B type TNBC) cells using trans-well assays (32). Consistent with our findings, Mitra et al. reported that knockdown of CPSF73 resulted in slower migration of fibroblasts, while increased expression of CPSF73 in fibroblast cells promoted their migration into the denuded area in a wound healing assay (78).

However, the extent of APA dynamics and molecular mechanisms that coordinate the phenotypic reduction of invasiveness has not been systematically elucidated. In our study, analysis of EMT-related proteins showed that JTE-607 effectively reverses the expression of EMT-related markers across all four tested cell lines. Our 3’ end-focused sequencing analysis of JTE-607-treated MDA-MB-231 revealed APA changes in mRNAs from many genes with well-characterized roles in promoting EMT. We further analyzed a subset of these that exhibited 3’ UTR lengthening and downregulation, including TGFβR1, SMAD4, TBX2, PAK1, CD44, ERBB2, and AKT2 and found that 3’ UTR lengthening in these genes was accompanied by decreased protein expression. These proteins mediate EMT via multiple signaling pathways.

An overview of the role of these genes in promoting EMT is provided in Supplementary Table 3. Briefly, activation of TGFβR1 via TGF-β stimulates Smad signaling to induce expression of Snail and ZEB and suppress E-cadherin (79). SMAD4 serves as a downstream effector of the canonical TGF-β signaling pathway, activating expression of EMT-promoting genes and suppressing the transcription of CDH1, which encodes E-cadherin (80). TBX2 is also a transcriptional repressor for CDH1 (81). PAK1 promotes EMT by facilitating cross-talk among multiple signaling pathways, including Ras, Wnt, and PI3K/AKT (82). CD44 regulates cell adhesion and activates multiple signaling pathways contributing to EMT and cancer progression (83). ERBB2 overexpression also enhances cell motility and pushes cancer cells toward EMT (84). While the shortening of PAK1 and CD44 has been observed in several cancer types compared to normal cells (21), to our knowledge, APA in the other candidate genes has not been reported. Together, these findings suggest that the production of longer mRNA isoforms encoding key drivers of EMT contributes to reversal of the EMT process and a reduction of invasion when CPSF73 is inactivated. Overall, our findings demonstrate that differential gene expression and APA changes directed by CPSF73 inhibition shape multiple inputs that regulate EMT and are central to the invasion and metastatic characteristics of cancer cells.

Our subsequent analysis of AKT2 indicated that these APA changes could indeed have functional consequences. We focused on AKT2 because of: (1) its well-documented promotion of migration and invasion through EMT activation, (2) significant 3’ UTR lengthening, with a RED value of 2.47 following JTE-607 treatment, and a corresponding decrease in Akt2 protein, and (3) the association of its high expression with shorter overall survival in breast cancer patients. Blocking use of the upstream PAS using AMOs induced 3’ UTR lengthening of AKT2 mRNA and the suppression of Akt2 protein and other EMT-related markers, including Snail1, N-cadherin, and Zeb1. Importantly, the invasion capacity in MDA-MB-231 cells was strongly reduced compared to the control. These results indicate that APA regulation of AKT2 alone can drive partial EMT reversal and suppress invasion. However, given the enrichment of crucial EMT-related genes among the ones exhibiting APA changes after CPSF73 inhibition, it is likely that modulating PAS choice in multiple mRNA transcripts, in addition to AKT2, contributes to the full EMT phenotype characteristics of cancer cells.

In terms of the functional consequences of different mRNA isoforms, extension of the 3’ UTR can create binding sites for regulatory elements such as microRNAs (miRNAs) and RNA-binding proteins (RBPs), which impact the stability, translation efficiency, and localization of the affected mRNAs (85). To try to identify RBPs that might influence 3′ UTR lengthening, we conducted a motif enrichment analysis alongside analysis of global protein/RNA crosslinking data, searching for predicted RBP binding sites within the regions 200 bp upstream and downstream of PASs, as well as in the region between the PASs of the seven genes whose APA was experimentally validated. While many RBPs were predicted to bind, we did not find a common RBP that might be regulating all or some of the tested genes. Similarly, in silico bioinformatics analysis of the sequence between the proximal and distal PASs using the microRNA prediction tool miRDB (86) did not identify any common miRNAs consistently binding to the sequence between PASs in the genes tested. Further research will be needed to determine the relevant RBPs and miRNAs and the functional consequences of their binding.

## Conclusion

Our findings identify APA as a mechanism regulating EMT and cancer cell invasion and show that chemical inhibition of CPSF73, which leads to changes in 3’ UTR length of multiple EMT-related genes, promotes EMT reversal. Further studies are required to fully elucidate the mechanisms by which APA influences EMT and metastasis in TNBC. The current study introduces a novel approach to reversing EMT by manipulating APA events. This goal could be accomplished generally by reducing the mRNA 3’ end processing capacity of the cell, as through the inhibition of CPSF73, or in a more focused way, by using antisense oligonucleotides to block the use of specific PASs whose APA changes may be drivers of EMT. A benefit to inhibiting CPSF73 to impair EMT-driven cancer cell invasion is the potential to simultaneously affect multiple signaling pathways at doses well tolerated by normal cells, while blocking a single APA event with AMOs offers specificity not always possible with activity-inhibiting drugs. Given that EMT, which is considered an early event in metastasis, is also implicated in the acquisition of therapeutic resistance and stem-like characteristics in cancer cells (87, 88), interfering with EMT-specific APA events may also prove beneficial in targeting properties that contribute to cancer relapse. Together, our results support a functional correlation between APA and the EMT process that could lead to a new way of suppressing cancer cells and blocking metastasis.

## Supporting information

Supplementary Table 1. List of antibodies used in this study.

Supplementary Table 2. List of primers used in this study.

Supplementary Table 3. Overview of the roles of genes with altered 3' UTRs following JTE-607 treatment in promoting EMT.

Supplemental File 1. The complete list of DEGs (MDA-DESeq2.JTE.v.CTRL)

Supplementary File 2. The most affected pathways following JTE-607 treatment based on GSEA analysis.

Supplementary File 3. 3' UTR APA changes. (MDA_APA_report_REDu (adjusted p value))

Supplementary File 4. Intronic APA changes. (MDA-APA_report_REDi (adjusted p value))

Supplementary File 5. List of EMT-related gene sets extracted from bioCarta-misgDB-GeneCard databases.

## Declarations

### Ethics approval and consent to participate

Not applicable.

### Consent for publication

All authors provide their consent to the publication of this work and have viewed and approved the final published version. This manuscript does not contain any individual person’s data in any form (including any individual details, images, or videos) and thus does not require any individual’s consent for publication.

### Availability of data and materials

The data generated in this study are available within the article and its supplementary data files. The sequence data generated in this study have been submitted to the Gene Expression Omnibus. Further information, requests for resources, and access to raw data should be directed to and will be fulfilled by the lead contact, Claire L. Moore.

### Author contributions

CM, MN, and HL conceived the study and designed the experiments. MN developed the strategy and methodology, performed the experiments, analyzed and reported the data, and organized the findings. ER performed and contributed to experimental assays. LW, SMM, and BT contributed to the computational analyses. MN wrote the first draft of the manuscript.

## Acknowledgements

We would like to acknowledge the Tufts Research Technology team for their advice and guidance, especially regarding the use of the Tufts University High-Performance Computing (HPC) Cluster for our analysis. We also thank all members of the Moore laboratory for their valuable input on this project and their comments on the manuscript.

---

**Supplementary Fig. 1.**
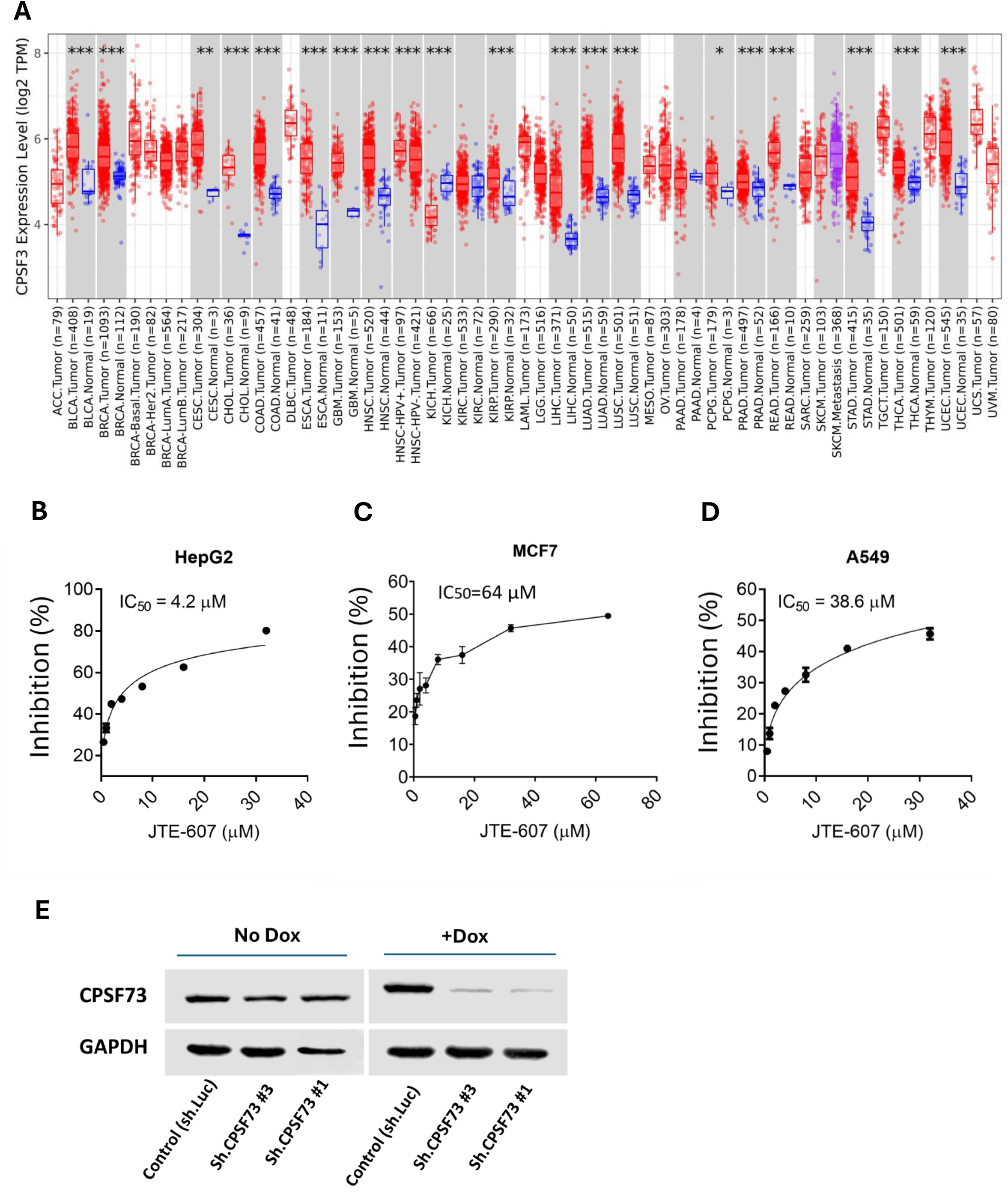
Analysis of CPSF73 expression in cancers and IC50 value determination of JTE-607 for cancer cell lines. **(A)** Expression of CPSF73 in different human cancers based on TIMER2.0 database analysis. Normal tissue is represented in blue, and tumor tissue in red. *, P<0.05; **, P<0.01; ***, P<0.001. **(B-D)** Calculation of the 50% inhibitory concentrations (IC50) of JTE-607 for **(B)** HepG2, **(C)** MCF7, and **(D)** A549 cells. Cells were treated with a range of JTE-607 concentrations from 0 to 64 µM for three days, and growth inhibition was measured by SRB assay. Values are presented relative to the growth observed in untreated cells, normalized to 100%. **(E)** Inducible CPSF73 KD in MDA-MB-231 cells was confirmed by Western blot.

**Supplementary Fig. 2.**
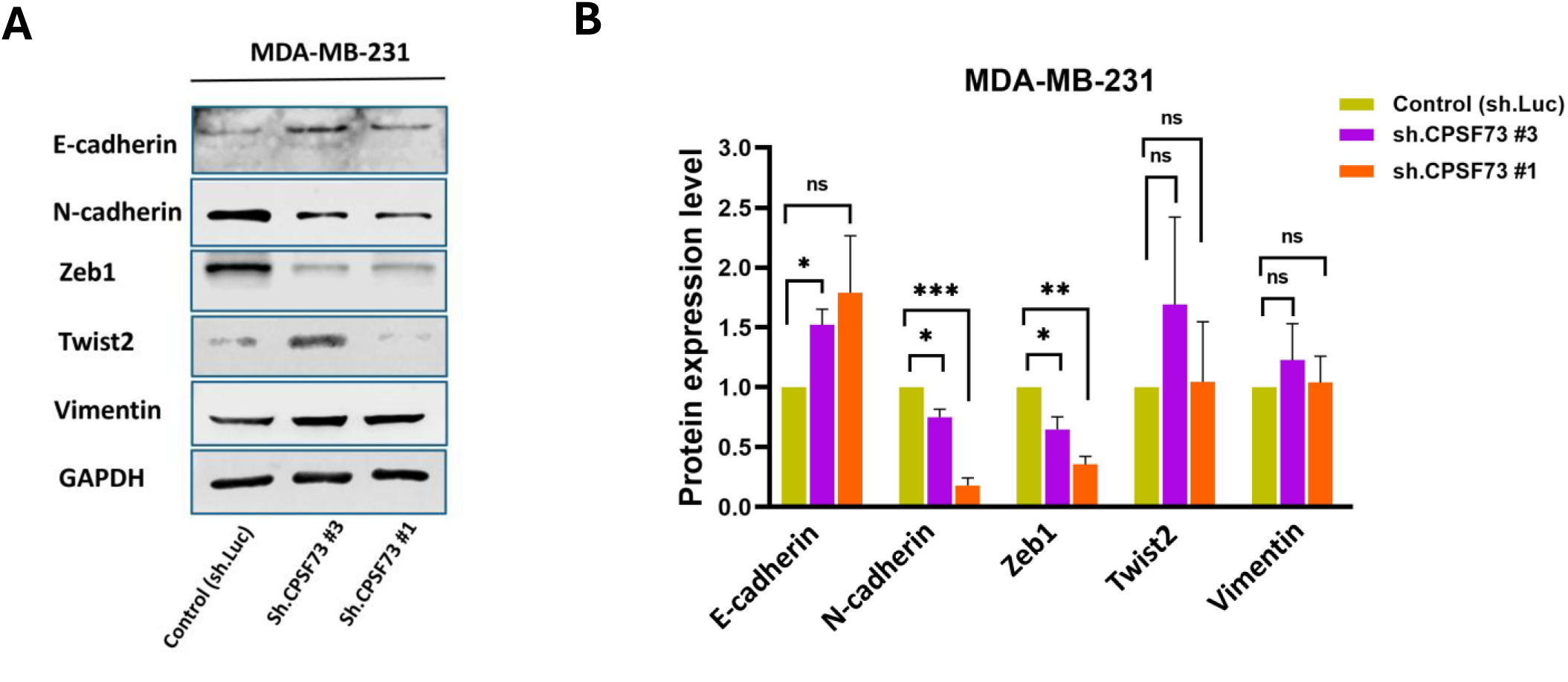
Western blot analysis demonstrating altered expression of EMT-related proteins in MDA-MB-231 cells with CPSF73 KD. **(A)** Representative western blot image and **(B)** corresponding densitometric bar graph quantifying the expression of EMT-related markers in MDA-MB-231 CPSF73 KD cells compared to the control. GAPDH was used to control the protein loading and normalize samples. Data are presented as mean ± SEM from at least three independent experiments (*p<.05, **p<.01, and ***p<.001).

**Supplementary Fig. 3.**
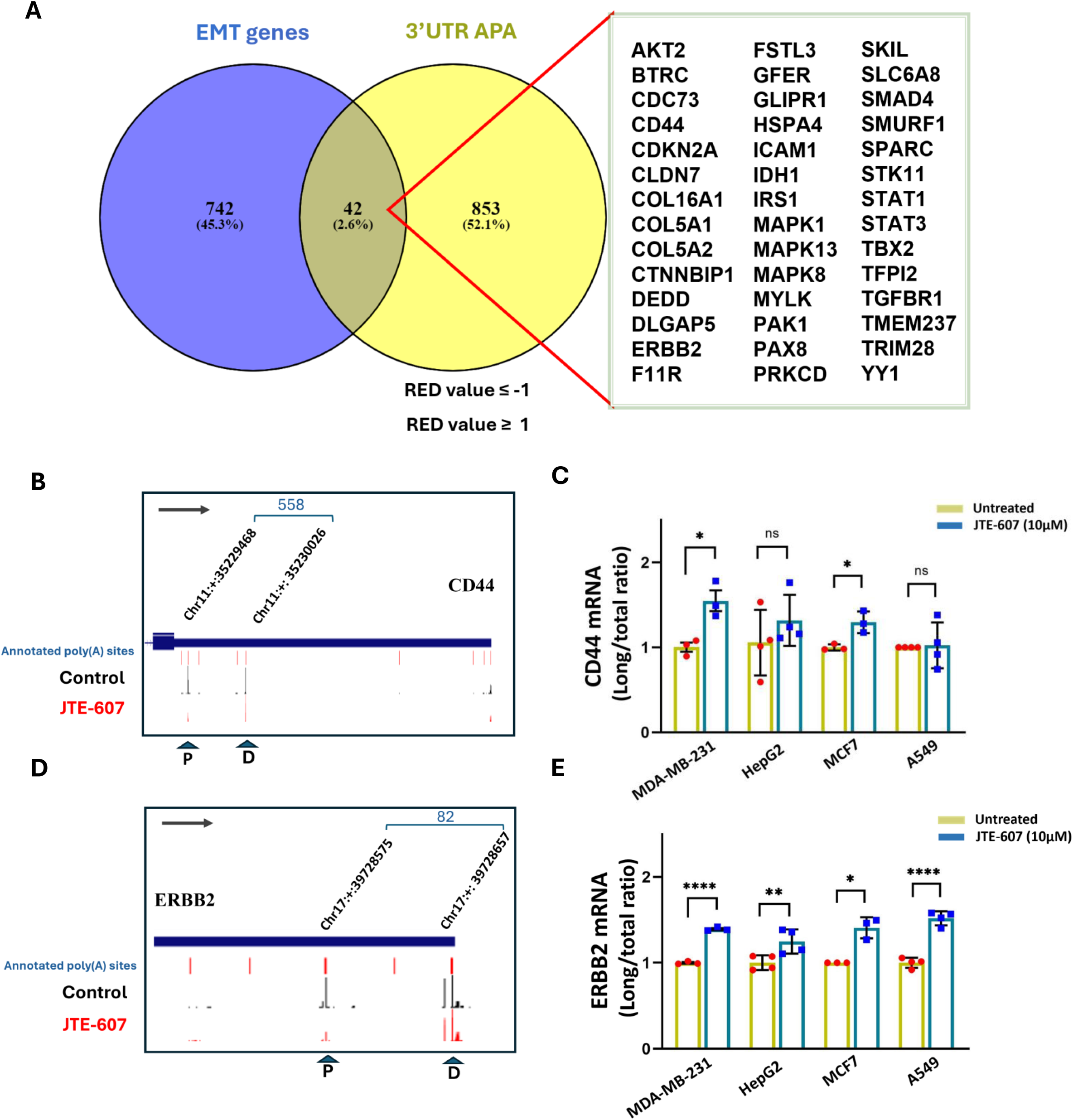
Venn diagram showing the intersection of genes with 3’ UTR APA changes and EMT-related genes. UCSC Genome Browser plots and RT-qPCR validation of selected EMT-related genes showing APA changes following CPSF73 inactivation. **(A)** The Venn diagram illustrates the intersection of EMT-related genes extracted from BioCarta, MSigDB, and GeneCard databases with all APA changes. **(B, D)** UCSC Genome Browser plots of RNA sequencing tracks showing changes in PAS usage, resulting in 3′ UTR profile alterations for CD44 and ERBB2 in MDA-MB-231 cells. Chromosome co-ordinates and distance between the differentially used PAS are indicated at the top of the plot. The positions of the distal (D) and proximal (P) PASs are indicated with arrows. The arrow inside the box shows the gene direction. **(C, E)** Corresponding bar graphs showing RT-qPCR validation of 3′ UTR profile alterations in all four cancer cell lines after CPSF73 inactivation using JTE-607 treatment. Gene expression levels were normalized to GAPDH. Data are presented as mean ± SEM from at least three independent experiments (*p<.05, **p<.01 and ***p<.001).

**Supplementary Fig. 4.**
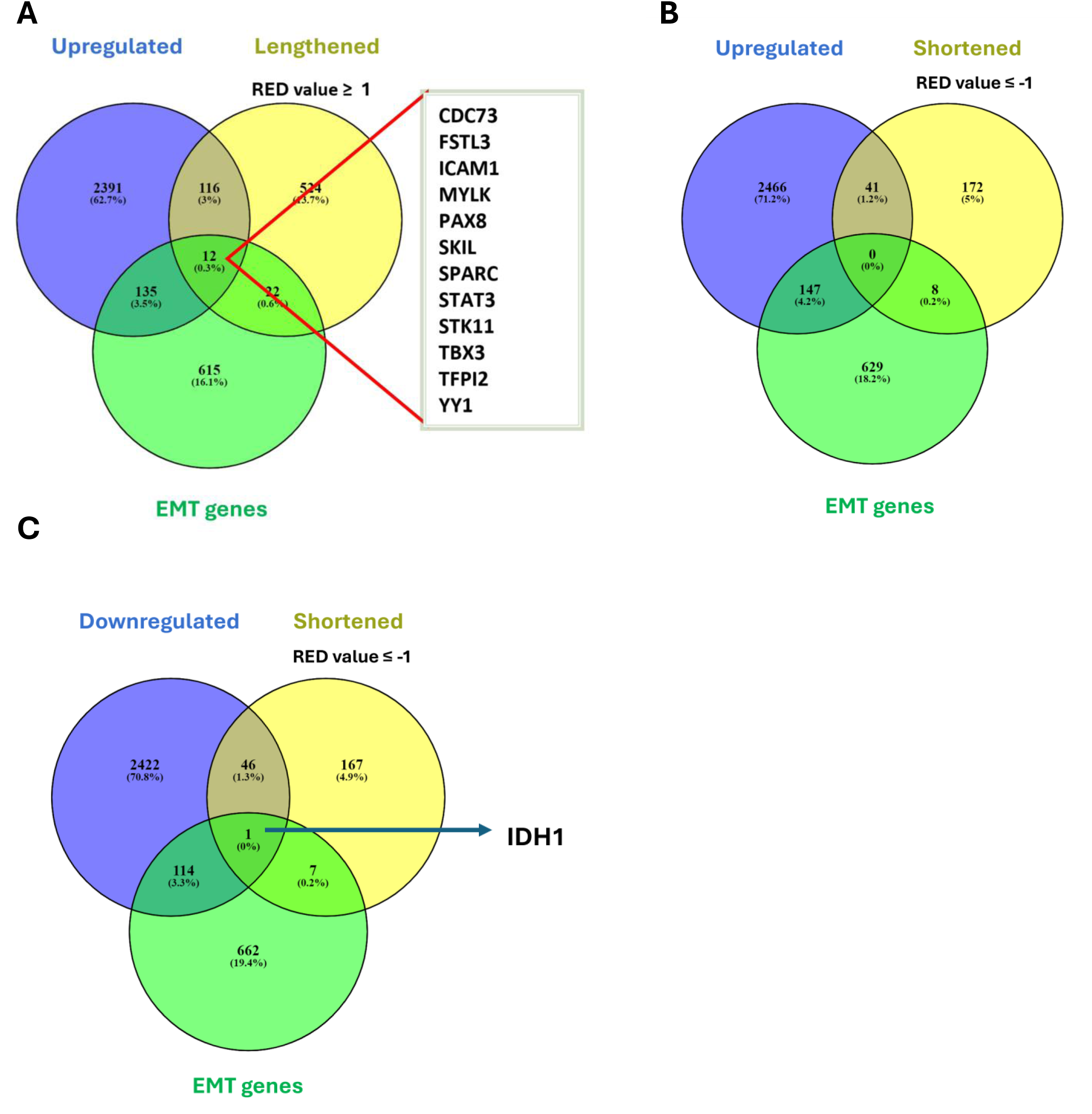
Venn diagram of genes showing 3’ UTR APA changes and their intersection with differentially expressed and EMT-related genes. The Venn diagram illustrates the intersection of EMT-related genes extracted from BioCarta, MSigDB, and GeneCard databases with **(A)** genes exhibiting both 3’ UTR lengthening and increased mRNA expression, **(B)** genes exhibiting both 3’ UTR shortening and increased mRNA expression, and **(C)** genes exhibiting both 3’ UTR shortening and decreased mRNA expression.

**Supplementary Fig. 5.**
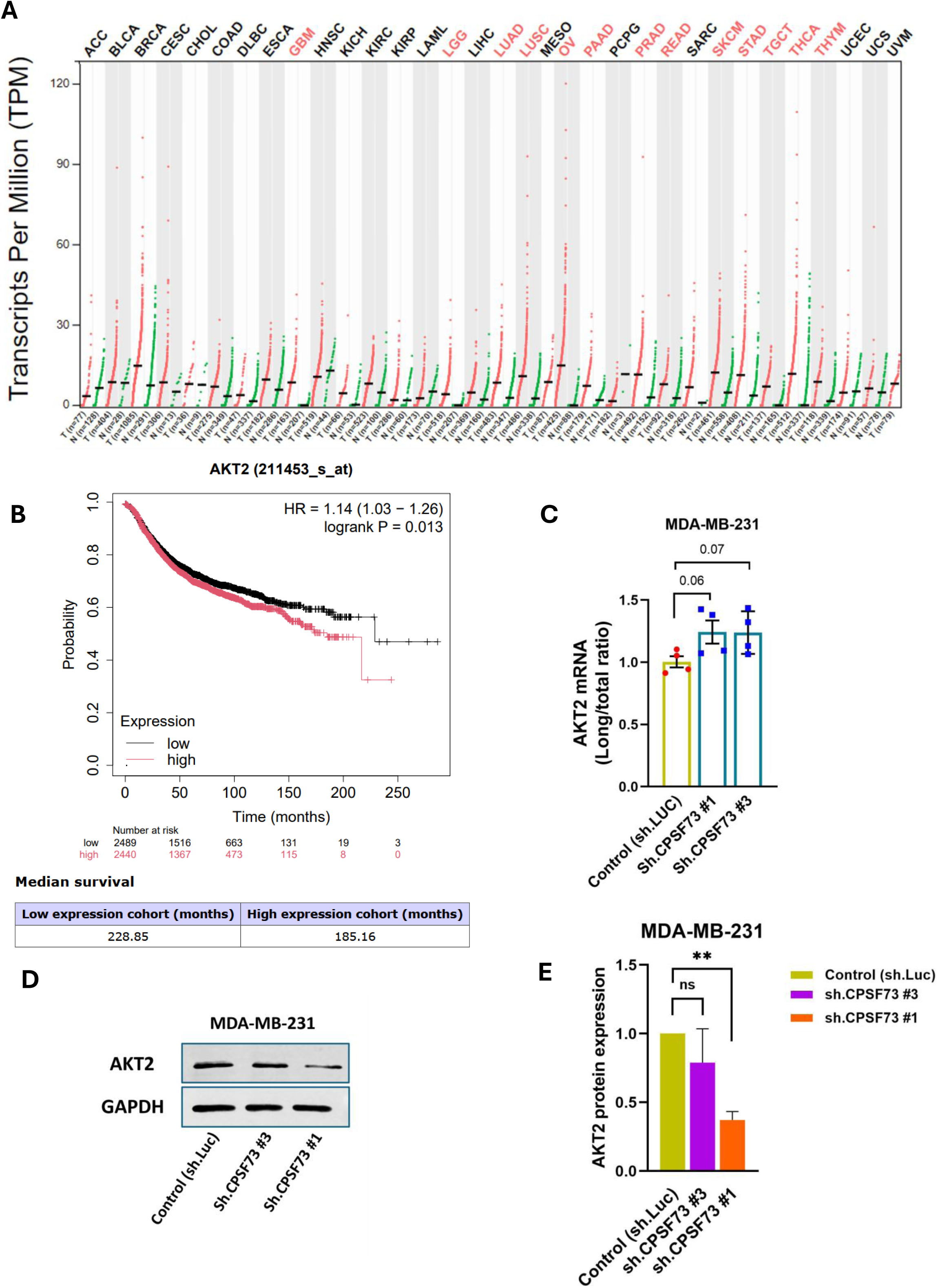
AKT2 is highly upregulated in breast tumor tissues and is associated with poor survival of cancer patients. **(A)** AKT2 transcripts are significantly upregulated in tumor tissues, including tissues from breast cancer, compared to normal tissues. Red and green dot plots represent AKT2 expression levels in tumor samples (red) and paired normal tissues (green), respectively. Data were retrieved from the GEPIA 2 database. **(B)** Kaplan-Meier survival analysis plot for AKT2 expression in breast invasive carcinoma (BRCA) shows the correlation between higher expression levels of AKT2 and poor outcomes in cancer patients. Data were retrieved from the Kaplan-Meier Plotter database. **(C)** RT-qPCR analysis of changes in the long/total ratio of AKT2 in MDA-MB-231 cells following CPSF73 KD. Gene expression levels were normalized to GAPDH. **(D** and **E)** Representative western blot image and corresponding densitometric bar graphs showing a significant decrease in the protein levels of Akt2 in MDA-MB-231cells with CPSF73 KD. GAPDH was used as a protein loading control for normalization. Data are presented as mean ± SEM from at least three independent experiments (**p<.01).

