## Supplementary Table 1. List of antibodies used in this study. for "Targeting CPSF73, the mRNA 3’ End Processing Endonuclease, Moves Cancer Cells Away from the Mesenchymal State"

| **Antibodies** | **SOURCE** | **IDENTIFIER** |
| --- | --- | --- |
| Mouse monoclonal anti-GAPDH antibody (G-9) | Santa Cruz Biotechnology | Cat# sc-365062; RRID: AB_10847862 |
| Mouse monoclonal anti-p21 Waf1/Cip1 antibody (F-5) | Santa Cruz Biotechnology | Cat# sc-6246; RRID: AB_628073 |
| Mouse monoclonal anti-N-cadherin antibody (13A9) | Santa Cruz Biotechnology | Cat# sc-59987; RRID: AB_781744 |
| Mouse monoclonal anti-Smad4 antibody (B-8) | Santa Cruz Biotechnology | Cat# sc-7966; RRID: AB_627905 |
| Mouse monoclonal anti-TBX2 antibody (D-3) | Santa Cruz Biotechnology | Cat# sc-514291; RRID: AB_2941848 |
| Mouse monoclonal anti-TGFβR1 antibody (D-1) | Santa Cruz Biotechnology | Cat# sc-518018 |
| Mouse monoclonal anti-SNAI1 antibody (E-10) | Santa Cruz Biotechnology | Cat# sc-393172; RRID: AB_2938534 |
| Mouse monoclonal anti-Akt2 antibody (1G8C12) | Santa Cruz Biotechnology | Cat# sc-81148; RRID: AB_1118811 |
| Mouse monoclonal anti-ZEB1 antibody (H-3) | Santa Cruz Biotechnology | Cat# sc-515797; RRID: AB_2934316 |
| Mouse monoclonal anti-twist antibody (Twist2C1a) | Santa Cruz Biotechnology | Cat# sc-81417; RRID: AB_1130910 |
| Mouse monoclonal anti-Vimentin antibody (E-5) | Santa Cruz Biotechnology | Cat# sc-373717; RRID: AB_10917747 |
| Mouse monoclonal anti-CPSF73 antibody (C-3) | Santa Cruz Biotechnology | Cat# sc-393001 |
| Mouse monoclonal anti-Myc antibody (9E10) | Santa Cruz Biotechnology | Cat# sc-40 |
| Rabbit polyclonal anti-DHRS2 antibody | Proteintech | Cat# 15735-1-AP; RRID: AB_2091847 |
| Rabbit polyclonal anti-PAK1 antibody | Cell Signaling | Cat# 2602T |
| Rat monoclonal anti-alpha Tubulin antibody (YOL1/34) | Thermo Fisher Scientific | Cat# MA1-80189 |
| Goat polyclonal anti-E-cadherin antibody | R&D systems | Cat# AF648 |
| Rabbit monoclonal anti-Akt2 antibody | Cell Signaling | Cat# 2964; RRID: AB_331162 |
| Goat anti-mouse HRP conjugate antibody | Bio-Rad | Cat# 1705047; RRID: AB_11125753 |
| Goat anti-rabbit (GAR)-HRP conjugate antibody | Bio-Rad | Cat# 1705046 |

**Table S1.** List of antibodies used in this study.
