## Supplementary Table 2. List of primers used in this study. for "Targeting CPSF73, the mRNA 3’ End Processing Endonuclease, Moves Cancer Cells Away from the Mesenchymal State"

**Table S2.** PCR primer sequences used in this study.

| **Gene name** | **name** | **sequence from 5’ to 3’** |
| --- | --- | --- |
| AKT2 total RT-qPCR primers | Akt2 T-F | CCAGTCCATCACAATCACACCC |
|  | Akt2 T-R | GCCTGAAGAAGAACTGGAAAGGG |
| AKT2 Long RT-qPCR primers | Akt2 L-F | CTGTGGGGCTGAGGGGTATG |
|  | Akt2 L-R | AGGCACCCTCCTCATTCTGG |
| TGFBR1 Total RT-qPCR primers | TGFBR1 T-F | GTG GCG GGG AGA AGA AGT TG |
|  | TGFBR1 T-R | ATC TCT GCC TCA CGG AAC CA |
| TGFBR1 Long RT-qPCR primers | TGFBR1 L-F | TAG GGG TGT GGG TCT TCC AT |
|  | TGFBR1 L-R | ATG TGC TGA CCC AAA GGC AA |
| SMAD4 Total RT-qPCR primers | SMAD4 T-F | CCA ACA TTC CTG TGG CTT CC |
|  | SMAD4 T-R | GCA GTC CTA CTT CCA GTC CAG |
| SMAD4 Long RT-qPCR primers | SMAD4 L-F | AAG TAA TGG CTC TGG GTT GGG |
|  | SMAD4 L-R | GCA AGG TAG CCA AGG TCA CT |
| TBX2 Total RT-qPCR primers | TBX2 T-F | CCCTTGGCTGCGGGAGA |
|  | TBX2 T-R | TGTCGTTTCGTTCTCGCTCC |
| TBX2 Long RT-qPCR primers | TBX2 L-F | GTG GCA CTT CTG CCT TGA GT |
|  | TBX2 L-R | TAC ACA GCA ATG TAC ACC TTT GAA T |
| PAK1 Total RT-qPCR primers | PAK1 T-F | ACC AGA TGT CAA CCT TGG GC |
|  | PAK1 T-R | GCA CAG ATA ACC CCA GAG CA |
| PAK1 Long RT-qPCR primers | PAK1 L-F | AGG CCA CCA AAT CCA TCC AG |
|  | PAK1 L-R | TTA GAG CAA GGC TGG GCA TC |
| CD44 Total RT-qPCR primers | CD44 T-F | CATCTACCCCAGCAACCCTA |
|  | CD44 T-R | CTGTCTGTGCTGTCGGTGAT |
| CD44 Long RT-qPCR primers | CD44 L-F | CGGGTGTGCTATGGATGGCT |
|  | CD44 L-R | TGGAAAGCAATGCCCAGGGA |
| ERBB2 Total RT-qPCR primers | ERBB2 T-F | CTC CCA CCT CTT GAT GCC AG |
|  | ERBB2 T-R | GAC ACC TAC GGC AGA GAA CC |
| ERBB2 Long RT-qPCR primers | ERBB2 L-F | CCT CCC CAT ACA ACA CCC AT |
|  | ERBB2 L-R | TGG TGT CAG TAT CCA GGC TTT |
| Myco RT-qPCR primers | Myco-F | GGCGAATGGGTGAGTAACACG |
|  | Myco-R | CGGATAACGCTTGCGACCTATG |
| MYC US RT-qPCR primers | MYC-US-F | ATTACAGGTGTGAGCCAGGG |
|  | MYC-US-R | AGCCTGCCTCTTTTCCACA |
| MYC Span RT-qPCR primers | MYC-Span-F | ATCATTGAGCCAAATCTTAAGTTGTG |
|  | MYC-Span-R | CTCTGAAGGGGCAATTGATGA |
| GAPDH RT-qPCR primers | GAPDH-F | TGATGACATCAAGAAGGTGGTGAAG |
|  | GAPDH-R | TCCTTGGAGGCCATGTGGGCCAT |
| RPL-13A RT-qPCR primers | RPL-13A_F | CAAGCGGATGAACACCAAC |
|  | RPL-13A_R | CGCTTTTTCTTGTCGTAGGG |
