## Supplementary Table 3. Overview of the roles of genes with altered 3' UTRs following JTE-607 treatment in promoting EMT. for "Targeting CPSF73, the mRNA 3’ End Processing Endonuclease, Moves Cancer Cells Away from the Mesenchymal State"

| **Gene name** | **Function in EMT and metastasis** | **References** |
| --- | --- | --- |
| **TGFβR1**  **(**Transforming growth factor-beta receptor type 1) | TGF-βR1transmits signals upon activation by TGF-β binding to its extracellular domain, which in turn activates both TGF-β Smad and non-Smad signaling pathways. Activation of TGF-β–Smad signaling triggers the EMT response in cells by inducing the expression of Snail and ZEB proteins and suppressing E-cadherin during the EMT process. | (86) |
| **SMAD4**  **(**SMAD family member 4) | SMAD4, a member of the Smad family of TFs, serves as a downstream effector and central signal transducer of the canonical TGF-β signaling pathway. It forms a complex with other SMADs (SMAD2/3) to facilitate their translocation to the nucleus, where it modulates the expression of EMT-related target genes. Moreover, SMAD4 acts as a transcriptional activator of FZD4, thereby promoting FZD4-dependent Wnt signaling. It also suppresses the transcription of the CDH1 gene, which encodes E-cadherin, by forming a complex with Snail1. While SMAD4 is associated with enhanced metastatic potential, SMAD4-deficient cancer cells tend to retain their epithelial characteristics. | (77, 87) |
| **TBX2**  (T-box transcription factor 2) | TBX2 belongs to the T-box gene family and is notably known to be a cell-autonomous inducer of EMT by acting as a direct transcriptional repressor of E-cadherin. Amplification and overexpression of TBX2 have been reported in aggressive human epithelial cancers. High expression levels of TBX2 are correlated with poor patient survival in colorectal, breast, pancreatic, and cervical cancers. Abrogation of TBX2 restores epithelial characteristics and abolishes tumor cell invasion in malignant human breast and colorectal cancer cells. | (78, 88) |
| **PAK1**  (The p21-activated Serine/Threonine kinase 1) | PAK1 is implicated in cancer progression and EMT, functioning as a critical node that stimulates and facilitates cross-talk among multiple signaling pathways, including Ras, Wnt, and PI3K/AKT. PAK1 is frequently upregulated in cancers, which promotes tumor growth and is thought to contribute to invasion and metastasis. | (79, 89) |
| **CD44** | CD44 is a transmembrane glycoprotein and a major component of the extracellular matrix (ECM) that functions as a receptor for hyaluronic acid (HA) and several components of the ECM, transducing signals to cytoskeletal proteins or the nucleus. The binding of HA to CD44 induces EMT. CD44 regulates cell-matrix adhesion, metastasis, and self-renewal, and it is also known as a CSC marker in many cancer types. | (80) |
| **ERBB2**  (Human Epidermal Growth Factor Receptor 2 (HER2)) | ERBB2, also known as Human Epidermal Growth Factor Receptor 2 (HER2), is a transmembrane tyrosine kinase receptor. In breast cancer, an increased expression of HER2 is associated with increased cell motility and metastasis. This overexpression of ERBB2 causes overactivation of downstream tyrosine kinase receptors, driving EMT through PI3K/Akt and MAPK pathways. | (81, 90) |
